# Istaroxime treatment ameliorates calcium dysregulation in a zebrafish model for Phospholamban R14del cardiomyopathy

**DOI:** 10.1101/2020.11.25.397422

**Authors:** S.M. Kamel, C.J.M. van Opbergen, C.D. Koopman, A.O. Verkerk, Y. L. Onderwater, S. Chocron, C. Polidoro Pontalti, M.A. Vos, T.P. de Boer, T.A.B. van Veen, J. Bakkers

**Affiliations:** Hubrecht Institute, Royal Netherlands Academy of Arts and Sciences (KNAW), University Medical Centre Utrecht, 3584 CT, Utrecht, The Netherlands; Department of Medical Physiology, Division of Heart & Lungs, University Medical Center Utrecht, Yalelaan 50, 3584 CM, Utrecht, The Netherlands; Department of Medical Biology, Amsterdam Cardiovascular Sciences, University of Amsterdam, Amsterdam University Medical Centers, Amsterdam, The Netherlands; Department of Experimental Cardiology, University of Amsterdam, Amsterdam University Medical Centers, Amsterdam, The Netherlands

**Keywords:** Phospholamban, PLN R14 mutation, zebrafish, arrhythmogenic cardiomyopathy, structural remodelling, cardiac electrophysiology, cardiac alternans, calcium dynamics, istaroxime

## Abstract

The heterozygous phospholamban (*PLN*) p.Arg14del (R14del) mutation is found in patients with dilated or arrhythmogenic cardiomyopathy. The PLN R14del mutation triggers cardiac contractile dysfunction and arrhythmogenesis by affecting intracellular Ca^2+^ dynamics. Little is known about the physiological processes preceding PLN R14del induced cardiomyopathy, which is characterized by sub-epicardial accumulation of fibrofatty tissue, and a specific drug treatment is currently lacking. Here, we addressed these issues using a knock-in PLN R14del zebrafish model. Hearts from adult zebrafish with the R14del mutation display age-related remodeling with sub-epicardial inflammation and fibrosis. Echocardiography revealed contractile pulsus alternans before overt structural changes occurred, which correlated at the cellular level with action potential duration (APD) alternans. These functional alterations are preceded by diminished Ca^2+^ transient amplitudes in embryonic hearts. We found that istaroxime treatment ameliorates the *in vivo* Ca^2+^ dysregulation, rescues the cellular APD alternans, while it improves cardiac relaxation. Thus, we present novel insight into the pathophysiology of *PLN* R14del cardiomyopathy and identify istaroxime as a potential novel drug for its treatment.

## Introduction

Arrhythmogenic cardiomyopathy (ACM) is a genetic inherited disease that occurs in 1:1000-1:5000 of the general population. ACM is characterized by the accumulation of fibrous tissue and fat at the sub-epicardial region of the ventricles where it infiltrates the myocardial wall. In addition, malignant ventricular arrhythmias and a high propensity for sudden cardiac death has been observed^1^. Sudden cardiac death most commonly occurs in young patients, during the concealed stage of the disease, when no overt cardiomyopathy is detectable yet. Little is known about the underlying cause of cardiac remodelling, eventually culminating in contractile dysfunction and heart failure, and sudden cardiac death in ACM patients^2^. Myocardial biopsies of late stage ACM patients also display inflammatory infiltrates, which are associated with the fibrofatty replacement of myocardial tissue^2^. Most cases of familial ACM are caused by mutations in desmosomal genes (e.g. plakophilin, desmoplakin). However, non-desmosomal gene mutations such as the phospholamban (*PLN*) p.Arg-14 deletion (R14del), have recently been identified as a cause for ACM as well^3^.

*PLN* R14del is a Dutch founder mutation and the most prevalent cardiomyopathy-related mutation in the Netherlands. It has been identified in 10-15% of all Dutch patients with dilated cardiomyopathy (DCM) and/or ACM and it is estimated that there are more than 1000 *PLN*-R14del Dutch carriers^4^. *PLN* mutations, including *PLN* R14del, have also been detected in several other countries, including Canada, the USA, Spain, Germany, and Greece^5–8^. Patient phenotypes are very heterogeneous, not only between families but also within families^9,10^. Up to today, no homozygous carriers have been identified^11^. Severely affected *PLN* R14del mutation carriers display profound fibrofatty infiltration in the sub-epicardial region of both ventricular walls, along with malignant ventricular arrhythmias and end stage heart failure^12^. Currently, specific drugs to treat this disease are lacking.

PLN is a small 52 amino-acid transmembrane sarcoplasmic reticulum (SR) protein and a crucial regulator of SR function within cardiomyocytes^13^. During excitation-contraction coupling, free cytosolic Ca^2+^ levels increase, causing more Ca^2+^ to bind to the cardiac myofilaments which generates contractile force. After contraction, free cytosolic Ca^2+^ levels must be diminished to resting levels in order to induce cardiomyocyte relaxation. Ca^2+^ is pumped back into the SR via the sarco(endo)plasmic reticulum Ca^2+^ ATPase (SERCA2a) and extruded out of the cell via the Na^+^-Ca^2+^ exchanger (NCX). Activity of SERCA2a, the cardiac SERCA isoform, is tightly regulated by its scaffolding protein PLN. Under physiological conditions, PLN inhibits SERCA2a activity and thereby tempers the amount of Ca^2+^ that flows back into the SR. Upon the inhibition of PLN activity, such as by phosphorylation on Ser^16^, SR Ca^2+^ cycling is enhanced leading to an increase in SR Ca^2+^ reuptake and improved cardiac function^14–16^. The *PLN* R14del mutation results in a protein that has a stronger affinity for SERCA2a^6^, which has been correlated with the disruption of the Ser^16^ phosphorylation motif. It was suggested that disruption of this motif could lead to an increased inhibition of the SERCA2a, cytoplasmic Ca^2+^ overload, and increased risk for malignant ventricular arrhythmias^17,18^. Whether changes in intracellular Ca^2+^ dynamics also occur *in vivo* and affect cardiac structure and function remains unaddressed. Studying these processes in depth requires accurate *in vivo* models.

During the past decades, the zebrafish (*Danio rerio*) has emerged as a powerful, cost-efficient, and easy-to-use vertebrate model to study human disease^19^. Its conserved genome (82% of all human disease causing genes has at least one zebrafish orthologue), physiology and pharmacology has made the zebrafish a highly valuable model for the resolution of human disease-related mechanisms and for novel drug (target) discovery^20–22^. In addition, CRISPR/Cas genome editing in the zebrafish has led to new opportunities, such as the introduction of patient specific gene mutations in the zebrafish genome^23–25^.

In this study, we used zebrafish with a R14del variant in the endogenous *plna* gene to understand the cardiac pathophysiology caused by the *PLN* R14del mutation. Strikingly, an age-related severe cardiac remodelling was observed at the sub-epicardial region of the heart, combined with a strong infiltration of immune cells, fibroblasts and fat deposits. In young-adult *plna* R14del zebrafish without apparent structural remodelling, we observed variations in ventricular outflow peak velocity between consecutive beats. In correlation with this, irregular action potential (AP) duration (APD) (alternans) were identified in isolated *plna* R14del cardiomyocytes. Since these irregular APs can be caused by altered Ca^2+^ dynamics, we analysed intracellular Ca^2+^ dynamics *in vivo*. Corroborating this we found that cardiac Ca^2+^ transient amplitudes were reduced in *plna* R14del hearts. Importantly, istaroxime ameliorated the *in vivo* Ca^2+^ dysregulation and contractile impairment in *plna* R14del zebrafish, and rescued the observed irregular APDs in *plna* R14del cardiomyocytes.

## Materials and Methods

### Zebrafish husbandry

Fish used in this study were housed under standard conditions as described previously^26^. All experiments were conducted in accordance with the ethical guidelines and approved by the local ethics committee of the Royal Dutch Academy of Sciences (KNAW).

### Generation of mutant lines

The R14del mutation was generated in the wild-type Tupfel Longfin (TL) strain zebrafish using CRISPR/Cas9 technology. *Plna* R14del fish were generated as described before^24^. In short, one-cell-stage zebrafish embryos were microinjected with an injection mixture consisting of (final concentrations): 150 ng/μl nuclear Cas9 (nCas9) mRNA, 20-40 ng/μl sgRNA, 10% (v/v) Phenol Red and 25 ng/μl template oligo. Each putative founder adult fish was crossed with a wild-type adult fish (F1). Homozygous fish (F2) were generated by inbreeding heterozygous mutant carriers.

### Adult zebrafish heart isolations and preparation

For paraffin sections, adult zebrafish hearts were dissected and fixed in 4% paraformaldehyde (dissolved in phosphate buffer containing 4% sucrose) at 4°C overnight, washed twice in PBS, dehydrated in EtOH, and embedded in paraffin. Serial sections were made at 10 μm using a microtome (Leica RM2035). For cryosections, zebrafish hearts were extracted and fixed in 4% paraformaldehyde (in Phosphate buffer) for 4 hours at room temperature (RT). Three washes of 30 minutes were performed using 4% sucrose (in Phosphate buffer) followed by an overnight incubation at 4°C in 30% sucrose (in Phosphate buffer). Hearts were embedded in tissue freezing medium (Leica, Lot# 03811456), frozen on dry ice and kept at −80°C. Cryo-sectioning using Cryostar NK70 (Thermo Scientific) was performed to obtain 10 μm thin sections. Images of extracted whole hearts were acquired using a Leica M165 FC stereo microscope.

### Adult zebrafish heart staining

In situ hybridization (ISH) was performed on paraffin sections as described previously^27^, with the exception that the hybridization buffer did not contain heparin and yeast total RNA. Primers for the *grn1* probe were: forward primer TCCCGGTGGAGACTGTAGAC, reverse primer AATGACGGTGCATTTTGACA. Primers for the *postnb* probe were: forward primer AGAGGTTCTGGACAGGCTCA, reverse primer AAGGCACCATTTTTCACCAG. The *myl7* and *tbx18* ISH probes were as described previously^28,29^. Hematoxylin and Eosin (H&E) staining was performed on cryosections according to standard laboratory protocol for paraffin sections. Cryosections were fixed in 4% Paraformaldehyde for 1 hour at RT beforehand. Pico-sirius Red staining on cryo-sections was performed in accordance with standard laboratory protocol. Acid fuchsin-orange (AFOG) and Oil Red O (ORO) staining were performed on cryosections as described previously^30,31^. Imaging of stained sections was performed using Leica DM4000 B LED upright automated microscope. TUNEL apoptosis staining was performed on cryosections and detected using In Situ Cell Death Detection Kit, fluorescein from Roche (Mannheim, Germany, Lot#29086800) according to manufacturer’s instructions. Nuclei were staining with DAPI (4′,6-diamidino-2-phnylindole) from Molecular Probes. Confocal images were acquired using a Leica Sp8 confocal microscope and processed using Imaris image analysis software (version 9.3.1).

### Echocardiography

Adult zebrafish were anesthetised in a cage filed with 4% MS-222 in aquarium water. When the zebrafish were unresponsive to a slight tail pinch with forceps it was determined that the appropriate anesthetised condition for echocardiography was reached. Anesthetized zebrafish were placed in a custom-made mold and placed ventral side up in a large petridish filled with 4% MS-222 in aquarium water. Breathing was monitored by visual tracking of opercular movement to ensure fish health throughout the protocol. Transthoracic echocardiography was performed using a Vevo2100 Imaging System (VisualSonics Inc., Toronto, Canada) with a 50 MHz ultrasound transducer fixed above the ventral side of the zebrafish and parallel to the longitudinal axis plane. Recordings were performed within 3 minutes after induction of anesthesia to preserve optimal cardiac performance. Quantitative measurements were assessed offline using the Vevo2100 analytical software. Color Doppler images were used for measuring heart rate, ventricular outflow diameter (manually), ventricular outflow tract velocity time integral (VOT VTI) and ventricular outflow peak velocity (VOT PV), as validated by Wang *et al.* and Visual Sonics Inc.^32^. Every parameter was determined as the average measurement of at least five cardiac cycles. Ventricular outflow surface was calculated by the equation π((OT diameter*OT diameter)/2), VOT diameter was measured in 3 consecutive beats per fish. Stroke volume was calculated by multiplying the VOT VTI with the ventricular outflow surface. Variation in outflow peak velocity was determined by the discrepancy in VOT PV between two consecutive heart beats and averaged over at least 6 beats per fish. VOT PV variation was corrected for the average total OT PV per fish. Hearts of these fish were then individually isolated and processed as frozen samples for cryosectioning.

### Cellular electrophysiology

#### Cell preparation

Single ventricular cardiomyocytes were isolated by an enzymatic dissociation procedure as described previously^33^. Ventricles from 3-4 adult fishes (6-7 months) were pooled and stored at RT in a modified Tyrode’s solution containing (in mmol/L): NaCl 140, KCl 5.4, CaCl_2_ 1.8, MgCl_2_ 1.0, glucose 5.5, HEPES 5.0; pH 7.4 (set with NaOH). Subsequently, the ventricles were cut in small pieces which were transferred to Tyrode’s solution with 10 μmol/L CaCl_2_ (30°C). The solution was refreshed one time before the addition of Liberase TM research grade (final concentration 0.038 mg/mL (Roche Diagnostics, GmbH, Mannheim, Germany)) and Elastase from porcine pancreas (final concentration 0.01 mg/mL (Bio-Connect B.V., Huissen, Netherlands)) for 12–15 minutes. During the incubation period, the tissue was triturated through a pipette (tip diameter: 2.0 mm). The dissociation was stopped by transferring the ventricular pieces into a modified Kraft-Brühe solution (30°C) containing (in mmol/L): KCl 85, K_2_HPO_4_ 30, MgSO_4_ 5.0, glucose 5.5, pyruvic acid 5.0, creatine 5.0, taurine 30, β-hydroxybutyric acid 5.0, succinic acid 5.0, BSA 1%, Na_2_ATP 2.0; pH 6.9 (set with KOH). The tissue pieces were triturated (pipette tip diameter: 0.8 mm) in Kraft-Brühe solution (30°C) for 4 min to obtain single cells. Finally, the cells were stored for at least 45 min in modified Kraft-Brühe solution before they were transferred into a recording chamber on the stage of an inverted microscope (Nikon Diaphot), and superfused with Tyrode’s solution (28°C). Quiescent single cells with smooth surfaces were selected for electrophysiological measurements.

#### Data acquisition

Action potentials (APs) and net membrane currents were recorded using the amphotericin-B perforated patch-clamp technique and an Axopatch 200B amplifier (Molecular Devices, Sunnyvale, CA, USA). Voltage control, data acquisition, and analysis were realized with custom-made software. Pipettes (resistance 3–4 MΩ) were pulled from borosilicate glass capillaries (Harvard Apparatus, UK) using a custom-made microelectrode puller, and filled with solution containing (in mmol/L): K-gluconate 125, KCl 20, NaCl 10, amphotericin-B 0.44, HEPES 10; pH 7.2 (set with KOH). Potentials were corrected for the calculated liquid junction potential ^34^. Signals were low-pass-filtered with a cut-off of 5 kHz and digitized at 40 and 5 kHz for APs and membrane currents, respectively. Cell membrane capacitance (C_m_) was estimated by dividing the time constant of the decay of the capacitive transient in response to 5 mV hyperpolarizing voltage clamp steps from –40 mV by the series resistance.

#### Action potentials

APs were elicited at 0.2 to 4 Hz by 3-ms, ~1.2X threshold current pulses through the patch pipette. Susceptibility to delayed afterdepolarization (DAD) generation was tested using fast burst pacing (20 APs at 3 Hz) which was followed by an 8-sec pause. After the 8-sec pause, a single AP was evoked to test the susceptibility of early afterdepolarizations (EADs) generation. APs were characterized by resting membrane potential (RMP), maximum AP amplitude (APA_max_), AP duration at 20, 50 and 90% of repolarization (APD_20_, APD_50_, APD_90_ respectively), maximal velocity (dV/dt) of the AP upstroke (Phase-0) and phase-3 repolarization (Phase-3), and plateau amplitude (APA_plat_; measured 50 ms after the AP upstroke). Averages were taken from 10 consecutive APs. DADs were defined as spontaneous depolarization of >1 mV.

#### Membrane currents

The AP measurements were alternated by a general voltage clamp protocol to elucidate the ionic mechanism underlying the AP changes. For K^+^ current measurements, 500 ms depolarizing and hyperpolarizing voltage clamp steps were applied from a holding potential of −50 mV with a cycle length of 2 s (Figure S3A). To ensure that the other cardiomyocytes in the recording chamber remained undistorted for biophysical analysis, our voltage clamp measurements were performed without specific channel blockers or modified solutions. Inward rectifier K^+^ current (I_K1_) and rapid delayed rectifier K^+^ current I_Kr_ were defined as the quasi steady-state current at the end of the voltage-clamp steps at potentials negative or positive to −30 mV, respectively. The L-type Ca^2+^ current (I_Ca,L_) was measured with a two-pulse voltage clamp protocol (Figure S3B) from a holding potential of −60 mV. The first pulse (P1) served to activate I_Ca,L_; the second pulse (P2) was used to analyse the inactivation properties of I_Ca,L_. I_Ca,L_ was defined as the difference between peak current and steady-state current. Current densities were obtained by normalizing to C_m_.

#### Istaroxime experiment

After baseline recordings, a number of isolated ventricular cardiomyocytes were treated for 5 minutes with 5 μM istaroxime (MedCham Exptress, Lot#11394). The effect on APD was measured at 1 and 4 Hz.

### High-speed brightfield imaging

Embryos were placed in 1-phenyl-2-thiourea (PTU) 20-24 hours post fertilization (hpf) to prevent pigmentation. 3 days post fertilization (dpf) embryos were embedded in 0.3% agarose prepared in E3 medium containing 16 mg/ml MS-222. Recordings were performed at 150 frames per seconds (fps) using a high-speed inverted light microscope at 28°C. Basal parameters were recorded first. Subsequently 3 mL of 100 μM istaroxime (MedCham Exptress, Lot#11394) or ouabain (Sigma-Aldrich, Lot#BCBZ9329) in E3-MS-222 solution was added, incubated for 30 minutes, and parameters were measured for a second time. Heart rate measurements and contractility parameters were analyzed using ImageJ (U. S. National Institutes of Health, Bethesda, Maryland, USA). Contraction time, relaxation time and contraction cycle time were analyzed by scrolling through the recorded movies frame by frame, and by identifying (1) the moment the ventricular wall moved inward and the ventricle started expelling blood (start of contraction), (2) the moment the ventricular wall moved back outward (start of relaxation), and (3) the moment that the ventricular wall reached its most dilated position (maximum relaxation). This process was repeated 3 times for each heart and values were averaged. The hemodynamic parameters such as surface area and volumes were analyzed using ImageJ by drawing an ellipse on top of the ventricle at end-diastole and end-systole. Per heart 6 ellipses were analyzed: 3 at diastole, and 3 at systole. Values were averaged. ImageJ provided the values for the minor and major axis of each ellipse. Surface area was calculated using the following formula: (0.5*major axis)*(0.5*minor axis)*π. End diastolic and end systolic volume (EDV/ESV) were calculated by: (1/6)*(π)*(major axis)*(minor axis2). Stroke volume (SV) by: EDV-ESV. Ejection fraction (EF) by: SV/EDV. Cardiac output (CO) by: SV*Heart rate.

### High-speed fluorescence imaging

*PLN* R14del fish were crossed to *tg*(*myl7*:Gal4FF; *UAS*:GCaMP6f) fish^35^ to obtain *PLN* R14del fish with a genetically encoded cardiac Ca^2+^ sensor. Wild-type fish, expressing GCaMP6f or a genetically encoded voltage sensor (VSFP Butterfly CY) were used as controls^35^. A morpholino (MO) oligomer targeted against *tnnt2a* (5′-CATGTTTGCTCTGATCTGACACGCA-3′) was injected at the 1-cell stage to uncouple contraction from excitation in embryos, thereby preventing motion artifacts in our recordings of intracellular cardiac Ca^2+^ handling. This ‘silent heart’ ATG morpholino was applied as described previously^36^. Embryos were placed in PTU after 20-24 hours to keep them transparent. 3 dpf embryos were embedded in 0.3% agarose prepared in E3 medium containing 16 mg/ml MS-222 and placed in a heated (28°C) recording chamber. Recordings were performed using a custom-build upright widefield microscope (Cairn research) equipped with a 20x 1.0 NA objective (Olympus XLUMPLFLN20X W). Blue LED excitation light (470 nm) was filtered using a 470/40 nm filter (Chroma ET470/40x) and reflected towards the objective using a 515 nm dichroic mirror (Chroma T515lp). Emitted fluorescence was filtered by a 514 long-pass filter (Semrock LP02-514RU) and images were projected on a high-speed camera (Andor Zyla 4.2 plus sCMOS). Recordings were performed at 100 fps, for 1000 frames. Basal parameters (heart rate, Ca^2+^ transient amplitude, diastolic Ca^2+^ level, upstroke time, recovery time) were recorded first. Subsequently, drug stocks were diluted in 28°C E3-MS-222 medium (istaroxime: MedCham Exptress, Lot#11394, ouabain: Sigma-aldrich, Lot#BCBZ9329) and the medium was mixed vigorously to assure a homogeneous concentration of the drug. Embryos were incubated for 30 minutes in the E3-MS-222 -istaroxime mixture and parameters were measured again. Recordings were analyzed using Image J and Matlab (Version R2015a, Mathworks).

### Quantitative PCR

RNA was isolated from hearts of adult TL wild-type fish. Adult hearts were separated into ventricular (n=3) and atrial (n=3) samples. *ef1α* was used as reference gene for the qPCR. Samples were loaded on 96-well PCR plate and run in CFX real-time PCR System (Bio rad) using KiCStart SYBR Green qPCR ReadyMix with ROX as recommended by the manufacturer (Sigma-Aldrich: KCQS02). Three technical replicates were performed for each sample together with no-template control (NTC) for each gene. The qPCR was performed with a 2-minute hold at 50°C and a 10 minute hot start at 95°C, followed by the amplification step for 43 cycles of 15 seconds denaturing at 95°C and 1 minute annealing/extension at 58°C, and the melting curve was obtained by increment of 1°C/ 10 seconds from 63°C to 95°C. Primers for qPCR were as follow: *plna* forward: TCTCCACTGCCATCTCTCCT and reverse: ACGAAGAGCTCCTGCATGTT, *plnb* forward: ACCAGCCTCATCATCTCCAC and reverse: ATCCTGTAGGTTGCGTTTGG, *ef1α* forward: CTTCTCAGGCTGACTGTGC and reverse: CCGCTAGCATTACCCTCC. Analysis was performed on the Cq values as described previously^37^. Fold Change was calculated using the formula 2^-(ΔΔCt).

### Statistical analysis

Statistical analysis and drawing of graphs and plots were carried out in GraphPad Prism (version 6 for Mac OS X and version 7 for Windows, GraphPad Software) and SigmaStat 3.5 software. Normality and equal variance assumptions were tested with the Kolmogorov-Smirnov and the Levene median test, respectively. Differences between two groups were analyzed using the paired Student’s t-test, comparisons between experimental groups were analyzed by one-way ANOVA for non-parametric variables with Tukey’s post-test for intergroup comparisons. Comparisons between experimental groups in combination with an intervention were analyzed by two-way ANOVA for non-parametric variables with Tukey’s post-test for intergroup comparisons. All data is presented as mean±SEM, and p<0.05 was considered significant. *p≤0.05, **p≤0.01, ***p≤0.001, ****p≤0.0001, n.s. p>0.05. N denotes the number of fish used per dataset.

## Results

### Cardiac remodeling in adult pln R14del zebrafish

A zebrafish homologue of human *PLN* has been annotated on chromosome 20 (ENSDARG00000069404), which we refer to as *plna* (**Figure S1A**). Subsequent screening for homologues sequences within the zebrafish genome revealed a second *pln*-like gene on chromosome 17 (si:ch211-270g19.5; ENSDARG00000097256), which we refer to as *plnb* (**Figure S1A**). Both zebrafish genes are expressed in the heart and have similar expression levels in the ventricle while in the atrium *plnb* is most abundant (**Figure S1C**). Both genes contain a short predicted open reading frame with high sequence similarity compared to human PLN (75% for Plna and 67% for Plnb) (**Figure S1B**). To get a better understanding of the consequences of the *PLN* R14del mutation, we engineered a fish line with a 3bp in-frame deletion in *plna* removing the conserved arginine at positions 14, which we will refer to as *plna* R14del (**Figure S1D**)^24^. Since the expression levels in the ventricle for *plna* and *plnb* are similar, we reasoned that zebrafish with a homozygous *plna* R14del mutation and wild type for the *plnb* gene (*plna* R14del/R14del; *plnb* +/+) would be the best genotype to represent the heterozygous PLN R14del mutation in patients. We will refer to this genotype as homozygous *plna* R14del fish or simply as *plna* R14del mutants. Adult fish with either a heterozygous or homozygous *plna* R14del mutation had a normal appearance and showed normal behaviour. To investigate cardiac morphological changes, we isolated hearts from 2-years-old wild type control, *plna* R14del heterozygous and homozygous zebrafish. Importantly, we observed that 23% of homozygous *plna* R14del fish displayed severe cardiac morphological changes, such as an increased size of the heart and a white flocculent layer of tissue that lined the outside of the heart. We did not observe any of these changes in wild-type siblings nor in heterozygous *plna* R14del fish (**Figure 1A-C**). Histological analysis of the remodelled hearts revealed altered tissue organisation in the sub-epicardial region with areas of high nuclear density and areas that appeared acellular compared to wild-type siblings and non-remodelled hearts (**Figure 1D-F**). Moreover, picro-sirius red staining revealed increased collagen deposition in the sub-epicardial region of these remodelled hearts, unlike wild-type siblings and non-remodelled hearts (**Figure 1G-I**). To address whether the morphological changes already occur earlier we isolated hearts from 10-month homozygous *plna* R14del fish (n=13) and observed one case with a similar altered tissue organisation, which was accompanied by cell death, in the sub-epicardial region (**Figure S2**). Interestingly, Oil Red O staining revealed prominent fat deposits throughout the sub-epicardial tissue and especially in the acellular areas (**Figure S2C&D**). Overall, these results suggest that zebrafish with a homozygous R14del mutation in *plna* and a wild type *plnb* gene undergo age-related cardiac remodeling most severe in the sub-epicardial region with incomplete penetrance.

**Figure 1:**
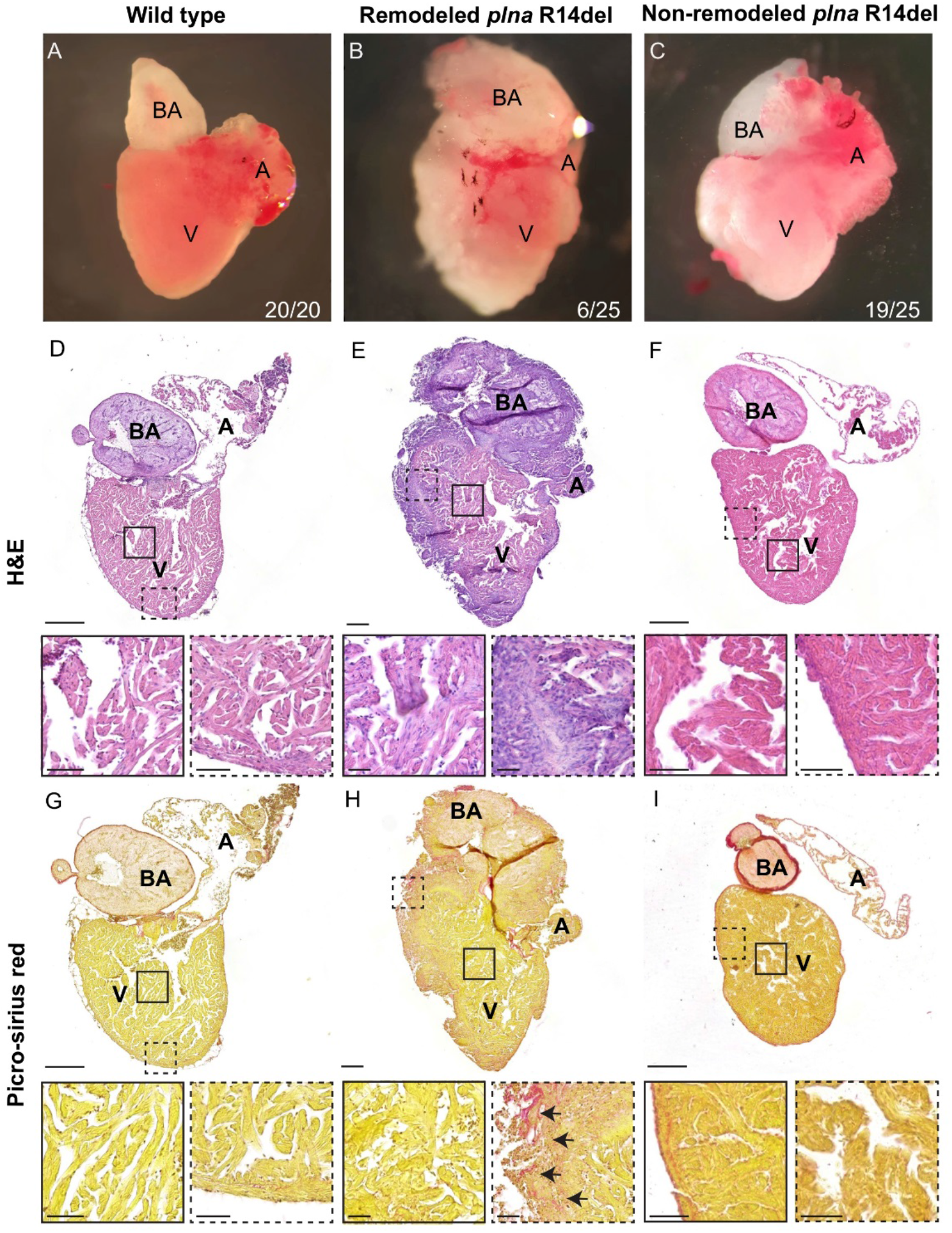
Structural remodelling of the adult *plna* R14del zebrafish heart. **A-C)** Bright field images of isolated adult zebrafish hearts, 2 years of age: wild-type fish, remodelled *plna* R14del mutant heart and non-remodelled *plna* R14del heart **D-F)** Haematoxylin and Eosin staining of the three conditions to identify nuclei, with zoom-in at indicated regions **G-I)** Picro-sirius red staining of collagen deposition for the three conditions, collagen fibers are shown as red staining. Zoom-in of each indicated region is included. Images were taken at a zoom of 20x. Scale bars are 200μm for whole heart tile scans and 50μm for zoom-in regions. A: atrium, V: ventricle, BA: bulbus arteriosus.

### Epicardial responses and immune infiltration in adult hearts of plna R14del zebrafish

Next, we aimed to identify which cell types could be involved in the observed cardiac remodeling and used *in situ* hybridization to investigate the presence of cell-type specific markers. T-box18 (Tbx18) is a transcription factor that is weakly expressed in the epicardium of the adult heart but its expression is induced upon cardiac injury^38^. Indeed, in wild-type hearts, we found expression of *tbx18* to be restricted to sparse cells in the sub-epicardial region (**Figure 2A**). Strikingly, expression of *tbx18* was clearly present in the epicardial region of *plna* R14del mutants, with *tbx18*-expressing cells forming several cell layers that covered the ventricle and the bulbus arteriosus (arrows in **Figure 2B**) and not overlapping with the myocardium, marked by *myl7* **(Figure 2 C &D)**. In addition, dispersed *tbx18* positive cells intermingled with cardiomyocytes were present in the ventricle of remodelled *plna* R14del mutant hearts, something we never observed in the hearts of wild types (**Fig. 2B**; boxes). Since TUNEL-staining revealed apoptotic cells in the sub-epicardial region we analysed the presence of immune cells by granulin (*grn1*) expression. Corroborating this finding, an accumulation of immune cells in the sub-epicardial region was observed as well (**Figure 2E&F**). As Picro-sirius red staining indicated enhanced fibrosis in the *plna* R14del mutant hearts, we analyzed the presence of activated fibroblasts, marked by the expression of periostin (*postnb*). Consistent with the increased fibrosis, we observed periostin expression in cells located in the sub-epicardial region (**Figure 2G&H).** Together these results indicate that cardiac remodeling of the sub-epicardial region in *plna* R14del mutants is accompanied by cellular damage, infiltrations of immune cells and the accumulation of fibroblasts.

**Figure 2:**
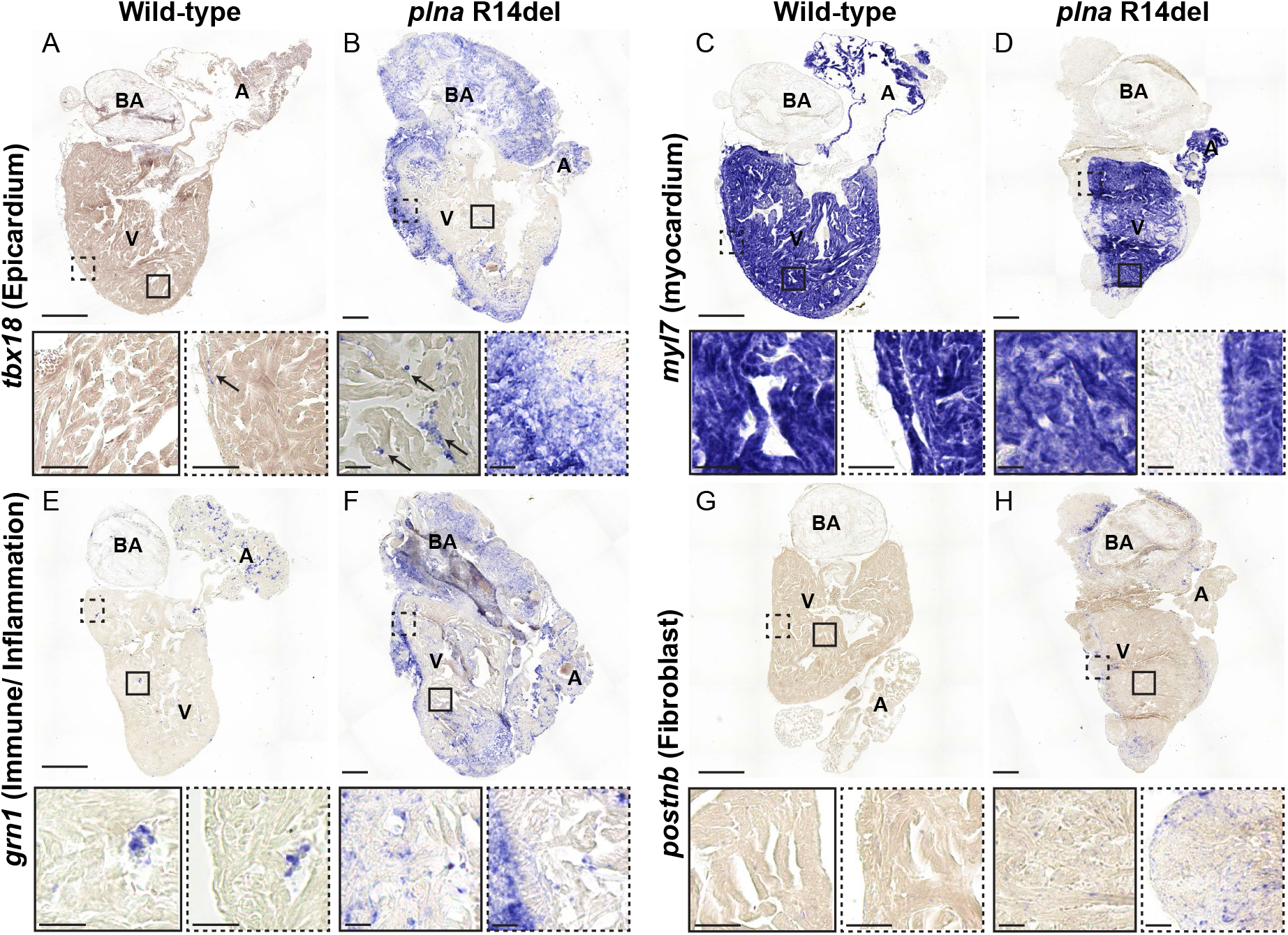
Epicardial response and immune infiltration in adult hearts of *plna* R14del zebrafish. *In situ* hybridisation of wild-type and *plna* R14del mutant adult zebrafish hearts, 2 years of age. **A-B)***tbx18* staining to indicate the epicardial cells, **C-D)***myl7* staining to indicate the myocardial cells, **E-F)***grn1* staining to indicate the immune cells, **G-H)***postnb* staining to indicate fibroblasts. Zoom-in of each region is indicated. Images were taken at a zoom of 20x. Scale bars are 200μm for whole heart tile scans and 50μm for zoom-in regions. A: atrium, V: ventricle, BA: bulbus arteriosus.

### Cardiac contractility defects in the plna R14del mutant heart

Structural remodeling of the heart is often preceded by impaired cardiac contractile dysfunction^39^. To examine cardiac contractility and pump function in *plna* R14del mutant zebrafish and relate this to heart morphology, we performed echocardiographic measurements combined with histological analysis of the cardiac tissue. For echocardiography fish were positioned ventral side up and a long-axis view, which included the two chambers of the heart, was imaged accordingly (**Figure 3A&B**). To track changes in cardiac performance over time, echocardiography of fish at two different ages (6 and 10 months old) was performed. Color doppler images were used to examine contractile parameters. While the heart rates were similar between groups, cardiac output was significantly higher in 10-month-old wild type and *plna* R14del mutant fish, compared to 6-month-old fish (**Figure 3C&D, Table S1**). Interestingly, *plna* R14del mutant fish showed alternations in ventricular outflow peak velocity (VOT PV) between consecutive beats, indicating alternating strong and weak beats. These so called pulsus alternans were already observed in 6-month-old *plna* R14del mutant fish, and became more apparent in 10-month-old fish resulting in a significantly higher extent of VOT PV variation in *plna* R14del mutant fish, compared to their wild-type siblings (**Figure 3E&F**). Histological analysis of hearts with strong pulsus alternans revealed no signs of cardiac remodelling (**Figure 3G**). Taken together, these results indicate that the *plna* R14del mutation causes beat-to-beat variations in cardiac output in absence of cardiac remodelling. This suggests that the cardiac contractile dysfunction is not caused by cardiac remodeling, but may be causal for cardiac remodelling.

**Figure 3.**
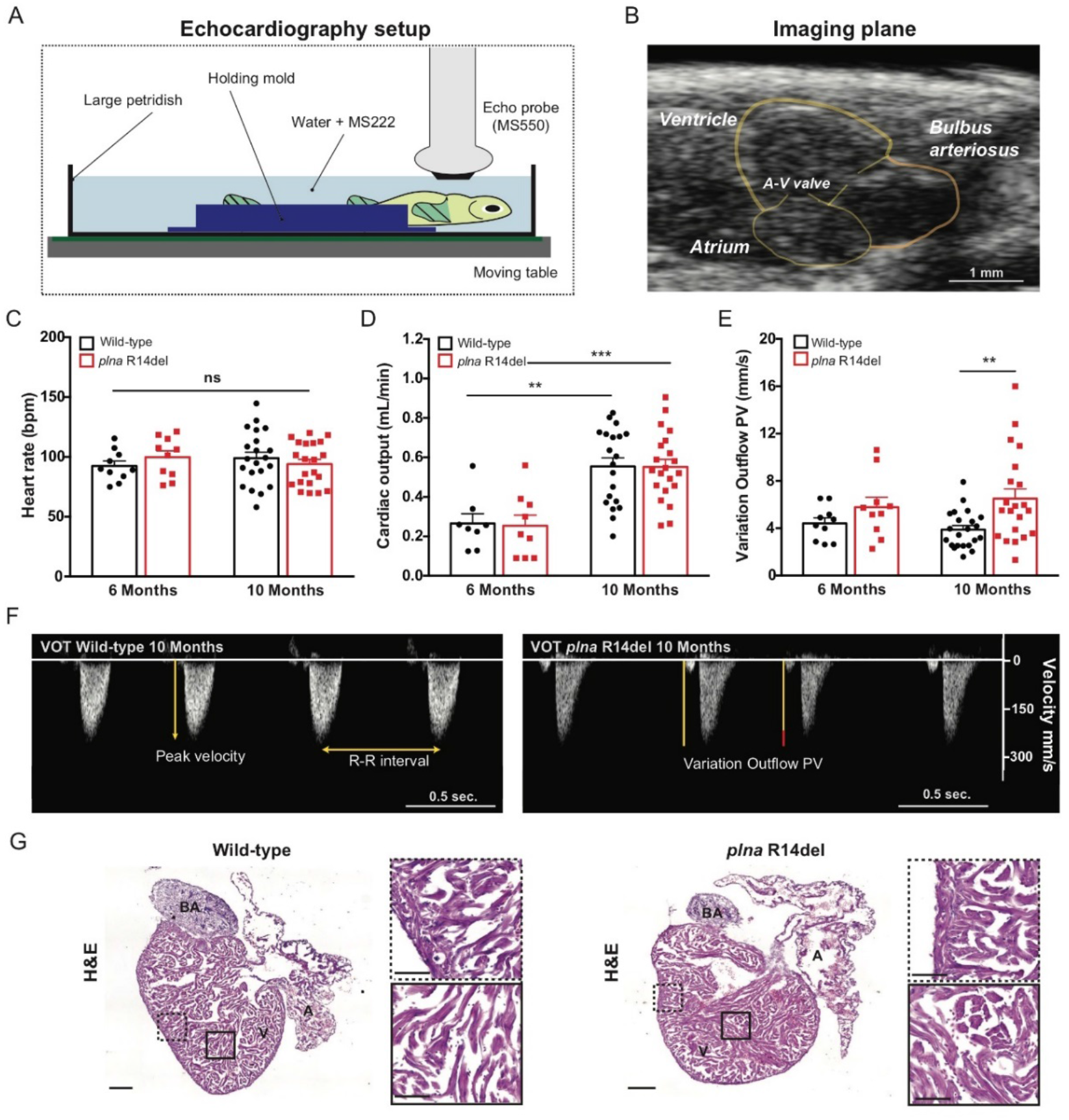
Contractile pulsus alternans in the structural normal adult *plna* R14del zebrafish heart. **A)** Graphical illustration of the zebrafish echocardiography imaging setup. **B)** B-mode echocardiography imaging plane of the adult zebrafish heart, ventricular walls and cardiac valves are depicted in yellow. **C)** Heart rate of wild-type (WT) and *plna* R14del mutant zebrafish, 6 and 10 months of age (mean ± SEM, WT n=10, *plna* R14del n=21, Two-way ANOVA). **D)** Cardiac output of WT and *plna* R14del zebrafish, 6 and 10 months of age (mean ± SEM, **p≤0.01, ***p≤0.001, WT n=10, *plna* R14del n=21, Two-way ANOVA). **E)** Variation in Outflow Peak Velocity in WT and *plna* R14del zebrafish, 6 and 10 months of age (mean ± SEM, **p≤0.01, WT n=10, *plna* R14del n=21, Two-way ANOVA). **F)** Representative examples of Colour Doppler ventricular outflow measurements in 10 months old WT and *plna* R14del zebrafish. **G)** Representative pictures of Hematoxylin & Eosin staining on 10 months old WT and *plna* R14del hearts (scale bar of 200μm for whole hearts as 50μm for zoom-in). Zoom-in of mid-myocardial and subepicardial region presented next to the corresponding whole heart picture. A: atrium, V: ventricle, BA: bulbus ateriosus, A-V valve: atrial-ventricular valve, Vent. Outflow: ventricular outflow, bpm: beats per minute, mL/min: millilitre per minute, mms/s: millimetre per second, VOT: ventricular outflow tract.

### Cellular electrophysiology of plna R14del ventricular cardiomyocytes

Cardiac (contractile) pulsus alternans can be the consequence of aberrancies in cardiac cellular electrophysiology and/or Ca^2+^ dynamics (reviewed by^40^). To examine cellular electrophysiology in *plna* R14del mutant zebrafish we performed patch clamp analysis on isolated cardiomyocytes from 10-month old fish. APs and membrane currents were measured using the perforated patch clamp technique, to limit technical disturbances in intracellular Ca^2+^ homeostasis. None of the AP parameters measured at 1 Hz differed significantly between WT and *plna* R14del mutant APs (**Figure 4A&B).** With increase in stimulus frequency, however, we observed a remarkable difference between both groups. While the average APD_90_-frequency relationships are virtually overlapping (**Figure 4C**), we noticed clear alternations in APD between consecutive APs (electrical alternans) in *plna* R14del mutant cells at higher pacing frequencies. **Figure 4D** shows a typical example of such an alternans. On average, the alternans in AP between consecutive beats was approximately 4-times larger in *plna* R14del mutant cells compared to cardiomyocytes from their wild-type siblings (**Figure 4E**).

**Figure 4.**
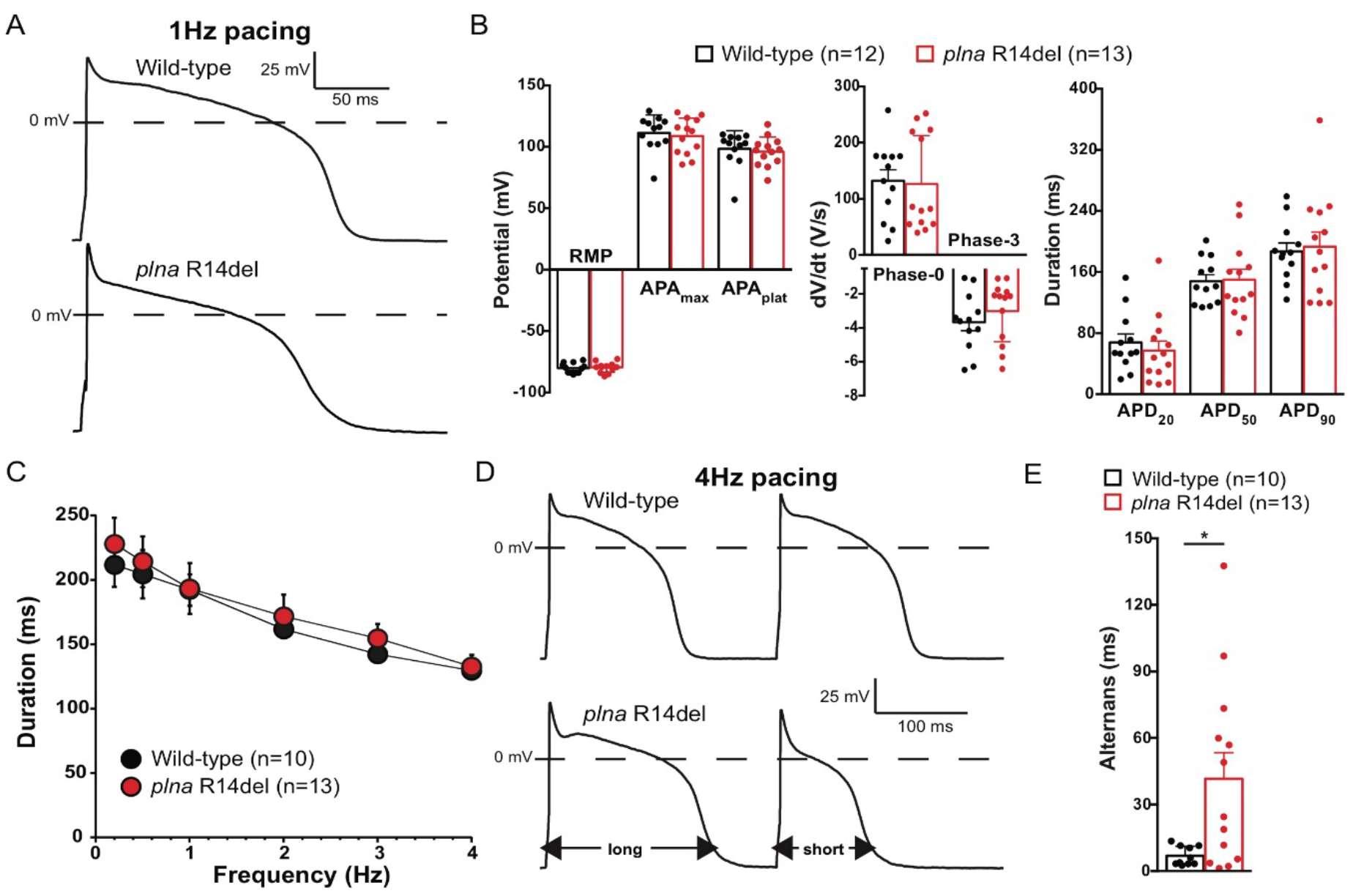
Action potential alternans in *plna* R14del isolated cardiomyocytes. **A)** Typical action potentials (APs) of cardiomyocytes isolated from a 10 months old wild-type (WT) (top panel) and *plna* R14del mutant (bottom panel) zebrafish, paced at 1 Hz. **B)** Average AP parameters at 1 Hz pacing (mean ± SEM, unpaired Students t-test). **C)** AP duration at 90% of repolarization (APD_90_) at 0.2 to 4 Hz pacing (mean±SEM, unpaired Students t-test). **D)** Two consecutive APs during 4 Hz pacing in a WT (top panel) and *plna* R14del (bottom panel) cardiomyocyte. The *plna* R14del cardiomyocyte showed an AP alternans where a long AP is followed by a short AP. **E)** Average APD_90_ difference between two consecutive APs (alternans) during 4 Hz pacing. WT: wild type, Hz: Hertz, mV: millivolt, ms: milliseconds, V/s: Volts per second, RMP: resting membrane potential, APA_max_: maximal action potential amplitude, APA_plat_: action potential amplitude at plateau phase, APD_20_: AP duration at 20% of repolarization, APD_50_: AP duration at 50% of repolarization, APD_90_: AP duration at 90% of repolarization.

We next measured the major membrane currents responsible for AP repolarization in zebrafish cardiomyocytes^41^. I_K1_, defined as steady-state current negative of −30 mV (**Figure S3A**), was not significantly different between WT and *plna* R14del mutant cardiomyocytes. This finding is in line with the comparable resting membrane potential (RMP) and Phase-3 repolarization velocities between the two groups **(Figure 4B).** In addition, I_Kr_ defined as steady-state current positive of −30 mV (**Figure S3A**), was unaffected in *plna* R14del mutant cardiomyocytes. Finally, we analysed I_Ca,L_ density and L-type Ca^2+^-channel (LTCC) gating properties. Neither current densities (**Figure S3B**) nor gating properties (data not shown) were affected by the *plna* R14del mutation. Together these data indicate that the *plna* R14del mutation does not affect the main ion channels underlying AP morphology.

### Impaired Ca^2+^ dynamics in plna R14del mutant hearts

AP alternans are frequently caused by a dysfunction of intracellular Ca^2+^ homeostasis and may trigger cardiac arrhythmias^42^. To study the effects of the *plna* R14del mutation on Ca^2+^ dynamics, we generated transgenic *plna* R14del zebrafish, expressing a cytosolic cardiac Ca^2+^ sensor (*tg*(*myl7*:Gal4FF *UAS*:GCaMP6f)). Analysis of GCaMP6f (cpEGFP) signal intensity over time allowed examination of Ca^2+^ transient amplitudes, diastolic Ca^2+^ levels and the speed of intracellular Ca^2+^ release and reuptake/clearance, as described earlier (**Figure 5A**)^35^. Here, we used 3-day old embryos since they allow *in vivo* analysis of intracellular Ca^2+^ dynamics due to their optical transparency. No significant differences were observed in Ca^2+^ transient frequency, upstroke time and recovery time between wild-type siblings and *plna* R14del mutants (**Figure 5B&C, S5**), indicating a normal speed of Ca^2+^ release into the cytosol and Ca^2+^ removal from the cytosol. However, the Ca^2+^ transient amplitude was on average 26% lower in *plna* R14del mutant embryos compared to wild-type siblings (WT: 99.8 ± 6.12% and *plna*-R14del: 73.9 ± 9.03, p≤0.05) (**Figure 5D**), suggesting that SR Ca^2+^ sequestration is impaired in the presence of the *plna* R14del mutation. Diastolic Ca^2+^ levels were not significantly affected in *pln*a R14del mutant fish (**Figure 5E**). Intracellular Ca^2+^ handling and cardiac contractility are very closely related processes, as free cytosolic Ca^2+^ ions bind troponin to cause actin-myosin linkages and contraction of the cardiomyocyte. High speed video imaging was used to analyze ventricular contractile parameters of the embryonic hearts (**Figure 5F**). No significant differences in stroke volume and cardiac output were observed between wild-type and *plna* R14del mutants (**Figure 5G&H).** Together these results indicate that intracellular Ca^2+^ dynamics of the embryonic heart are affected by the *plna* R14del mutation, but this has no consequences for the cardiac contractility at these early stages.

**Figure 5.**
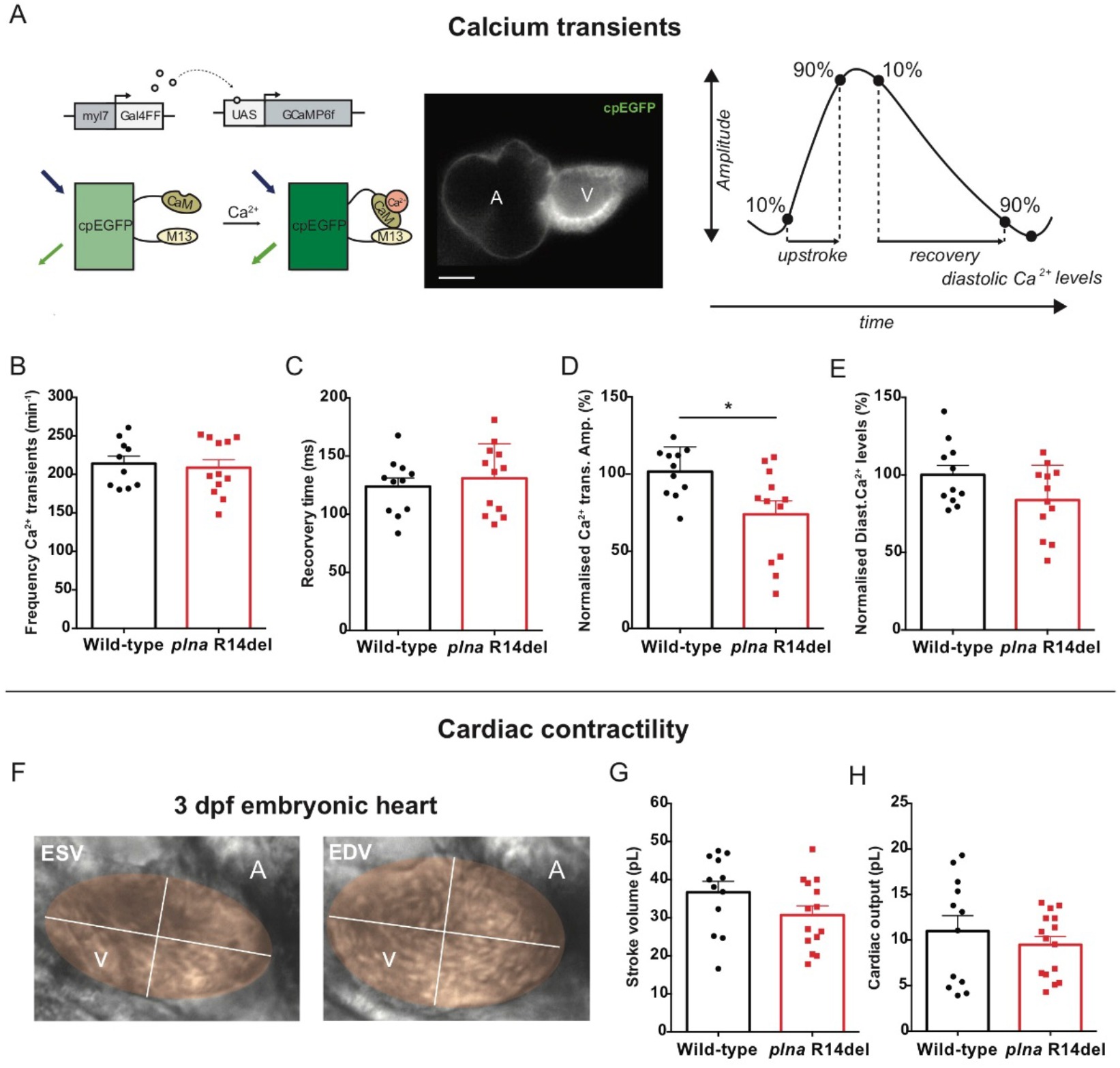
Reduced calcium transient amplitude in *plna* R14del embryonic zebrafish. **A)** DNA construct and sensor dynamics of GCaMP6f (left panel). GCaMP6f was placed under the control of the *myl7* promoter to restrict its expression to the heart. The *Gal4FF-UAS* system amplifies its expression. GCaMP6f consists of a circularly permutated enhanced green fluorescence protein (cpEGFP) fused to calmodulin (CaM) and the M13 peptide. When intracellular Calcium (Ca^2+^) rises, CaM binds to M13, causing increased brightness of cpEGFP. Using a high-speed epifluorescence microscope, movies of 3 dpf non-contracting GCaMP6f embryonic hearts were recorded (middle panel). Schematic representation of Ca^2+^ transient parameters examined from the recorded GCaMP6f signals (right panel). **B)** Frequency of Ca^2+^ transients, **C)** Ca^2+^ transient recovery time, **D)** normalized diastolic Ca^2+^ levels and **E)** normalized Ca^2+^ transient amplitude in wild-type (WT) and *plna* R14del embryonic zebrafish (mean ± SEM, *p≤0.05, WT n=10, *plna* R14del n=12, unpaired Students t-test). **F)** To measure hemodynamic parameters in high speed imaging recordings, we used ImageJ to fit an ellipse over the ventricle in every recording, both at end-systole (left) and end-diastole (right). Stroke volume **(G)** and cardiac **(H)** output in WT and *plna* R14del embryonic zebrafish (mean ± SEM, WT n=12, *plna* R14del n=14, unpaired Students t-test). CaM: calmodulin, UAS: upstream activation sequence: Ca^2+^: calcium, ESV: end systolic volume, EDV: end diastolic volume, bpm: beats per minute, ms: milliseconds, pL: Pico litre.

### Istaroxime enhances cardiac output by improving ventricular relaxation via targeting PLN-SERCA interaction

Since specific drug treatment is lacking to treat patients with PLN R14Del cardiomyopathy, we looked for a small molecule with the potential to rescue the above described electrophysiological phenotypes observed in *plna* R14del mutants. Istaroxime is known to stimulate the activity of SERCA2a, thereby enhancing SR Ca^2+^ sequestration^43^. In addition, it inhibits Na^+^/K^+^ -transporting adenosine triphosphatase (NKA) activity^44,45^. Since our results indicate that SR Ca^2+^ sequestration is impaired in *plna* R14del mutant zebrafish, we tested whether the treatment of *plna* R14del mutant embryos with istaroxime could improve the observed depression in Ca^2+^ transient amplitude. Wild-type and *plna* R14del mutant embryos with GCaMP6f were first imaged to record baseline Ca^2+^ dynamics. After baseline recording, 100 μM istaroxime was added to the E3-MS-222 medium and after 30 minutes of incubation another recording was performed (**Figure 6A**). 100 μM istaroxime was determined as the optimal drug concentration in a dose-response experiment using wild-type GCaMP6f fish (**Figure S4**). When combining the recordings of the GCaMP6f intensities from individual embryos before and after istaroxime treatment, we observed a near-complete restoration of the Ca^2+^ transient amplitude in *plna* R14del mutants, approaching wild-type levels (**Figure 6B&C)**, leaving diastolic Ca^2+^ levels unaffected (**Figure 6D**).

**Figure 6.**
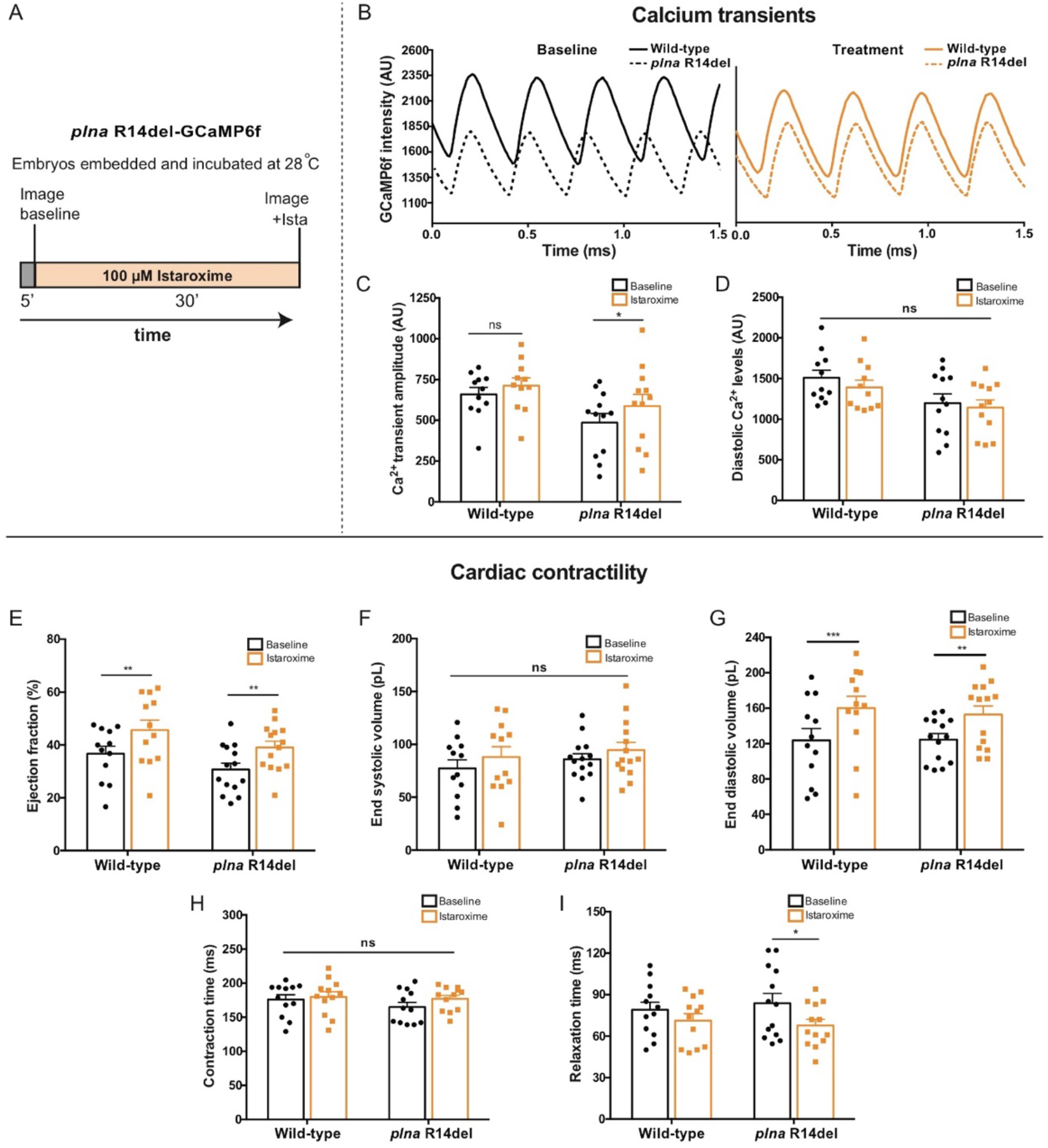
Istaroxime improves cardiac calcium dynamics and cardiac relaxation in *plna* R14del embryonic zebrafish. **A)** Overview of the experimental setup. Embryos were embedded in agarose and incubated for 5 minutes at 28°C. Baseline measurements were performed and subsequently embryos were incubated with 100 μM istaroxime for 30 minutes, and imaged again. **B)** Representative calcium (Ca^2+^) transients from one wild-type (WT) and one *plna* R14del embryonic zebrafish, before and after incubation with 100 μM istaroxime. **C)** Ca^2+^ transient amplitude and, **D)** diastolic Ca^2+^ levels in WT and *plna* R14del zebrafish, at baseline and after incubation with 100 μM istaroxime (mean ± SEM, *p≤0.05, WT baseline n=11, WT istaroxime n=11, *plna* R14del baseline n=12, *plna* R14del istaroxime n=13, Two-way ANOVA). **E-I)** Bar graphs of contractility parameters at baseline and after incubation with 100 μM istaroxime (mean ± SEM, *p≤0.05, **p≤0.01, ***p≤0.001, WT baseline n=12, WT istaroxime n=12, *plna* R14del baseline n=14, *plna* R14del istaroxime n=15, Two-way ANOVA), including end systolic volume (**E**), end diastolic volume (**F**), ejection fraction (**G**), contractile cycle length contraction time (**H**) and contractile cycle length relaxation time (**I**). AU: arbitrary units, ms: milliseconds, pL: Pico litre.

Istaroxime has shown to improve the systolic and diastolic function in acute heart failure patients^46,47^. In that regard, we examined the effect of istaroxime on cardiac contractility of wild-type and *plna* R14del mutants using high speed video imaging. As expected, istaroxime significantly enhanced stroke volume, cardiac output and ejection fraction in both wild-type and *plna* R14del mutants (**Figure 6E &S5**), which was not observed with a placebo treatment (**Figure S6)**. Stroke volume and cardiac output were likely affected via an improvement of the diastolic filling of the heart, as only end diastolic, and not end systolic volume, was changed by istaroxime treatment (**Figure 6F&G**). In order to further elucidate the effect of istaroxime on the rate of myocardial relaxation (lusitropy), we examined contractile cycle length. The total duration of contraction was divided into contraction time (time from maximal relaxation to maximal contraction) and relaxation time (time from maximal contraction to maximal relaxation). Total contraction duration and contraction time were not different between *plna* R14del mutants and wild-type embryos and were not affected by istaroxime treatment (**Figure 6H, S6**). Interestingly, the relaxation time was slightly longer in *plna* R14del mutants at baseline and significantly decreased after istaroxime treatment (**Figure 6I**).

Since istaroxime is known to block NKA and potentiates SERCA2a activity we tested whether istaroxime improves the Ca^2+^ transient amplitude via SERCA2a activation or NKA inhibition. Therefore, we compared the effect of istaroxime and ouabain, a specific NKA blocker, on ventricular Ca^2+^ transient dynamics in wild-type embryos. In contrast to istaroxime, ouabain treatment had no effect on Ca^2+^ transient amplitude (**Figure S7A&B**). To test whether a NKA block could be responsible for the lusitropic effects of istaroxime, potential changes in contractility were examined in ouabain treated embryos. Ouabain did not significantly enhance stroke volume, ejection fraction or cardiac output (**Figure S7C-F**). Next we validated the effectiveness of ouabain in our zebrafish model using embryos expressing a genetically encoded Voltage Sensitive Fluorescent Protein (VSFP Butterfly-CY) (**Figure S8A**). Fluorescent high-speed recordings of VSFP embryos demonstrated a clear APD shortening in the presence of ouabain compared to baseline values (**Figure S8B**).

Together, these data indicate that istaroxime can rescue impaired Ca^2+^ transient amplitudes in *plna* R14del mutant cardiomyocytes, an effect that is predominantly asserted via interference of the PLN-SERCA interaction.

### Istaroxime shortens repolarization and restores APD alternans in plna R14del cardiomyocytes

To test whether istaroxime could also rescue the AP alternans that we observed in adult *plna* R14del cardiomyocytes, we performed patch clamp recordings on isolated ventricular cardiomyocytes. To this end, cardiomyocytes of adult wild-type and *plna* R14del zebrafish hearts were measured at 1 and 4 Hz before and after treatment with istaroxime. Interestingly, istaroxime treatment resulted in a significant shortening of APD_20_, APD_50_ and APD_90_ and a complete absence of APD alternans (**Figure 7A-D**). The effects of istaroxime on APD were confirmed *in vivo*, using high-speed fluorescence imaging on embryos expressing VSFP Butterfly-CY (**Figure S8C**).

**Figure 7.**
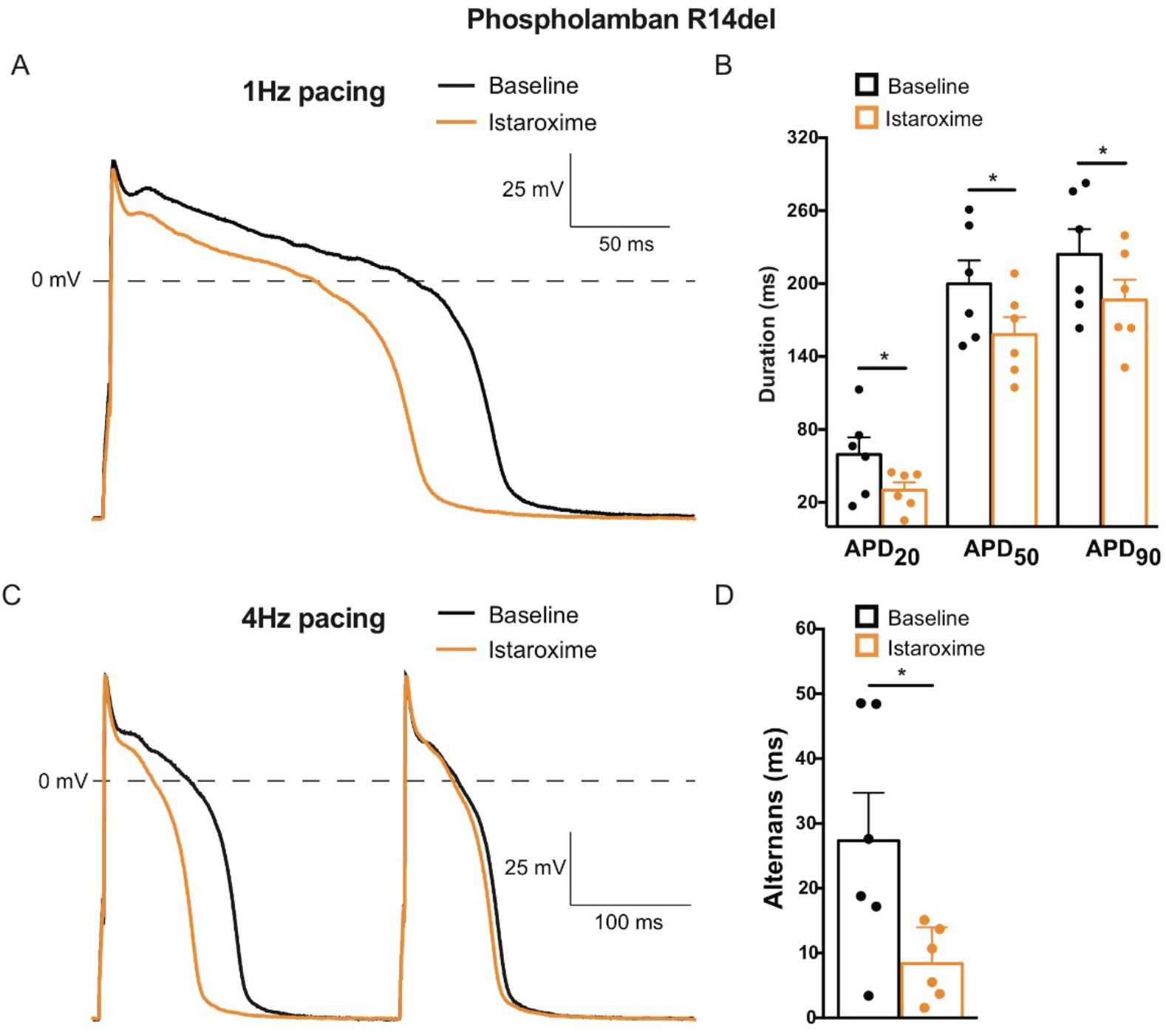
Istaroxime reverses action potential alternans in *plna* R14del isolated adult cardiomyocytes. **A)** Typical action potentials (APs) of *plna* R14del isolated adult cardiomyocytes at baseline (black line) and after incubation with 5 μM istaroxime (orange line), paced at 1 Hertz (Hz). **B)** Average AP parameters of *plna* R14del isolated adult cardiomyocytes at baseline (black bars) and after incubation with 5 μM istaroxime (orange bars), at 1 Hz pacing (mean ± SEM, *p≤0.01, baseline n=6, istaroxime n=6, unpaired Students t-test). **C)** Typical action potentials (APs) of *plna* R14del isolated adult cardiomyocytes at baseline (black line) and after incubation with 5 μM istaroxime (orange line), paced at 4 Hertz (Hz). The *plna* R14del cardiomyocyte showed an AP alternans at baseline, where a long AP is followed by a short AP. **D)** Average APD_90_ difference between two consecutive APs (alternans) at baseline and after incubation with 5 μM istaroxime, during 4 Hz pacing pacing (mean ± SEM, *p≤0.01, baseline n=6, istaroxime n=6, unpaired Students t-test). Hz: Hertz, mV: millivolt, APD_20_: AP duration at 20% of repolarization, APD_50_: AP duration at 50% of repolarization, APD_90_: AP duration at 90% of repolarization.

Together these results demonstrate that istaroxime rescues the APD alternans in *plna* R14del mutant cardiomyocytes and indicates that these APD alternans are caused by defects in intracellular Ca^2+^ handling.

## Discussion

In this study, we utilized the zebrafish to investigate the *in vivo* consequence of a PLN R14del mutation. Our results indicate that *plna* R14del results in alterations of cardiac intracellular Ca^2+^ dynamics already at the embryonic stages, and that these changes relate to a reduction in SR Ca^2+^ re-uptake. At this early age, the observed defects in Ca^2+^ handling do not lead to measurable differences in cardiac output. In the adult heart *plna* R14del cardiomyocytes start to show alterations in APD, especially under high pacing frequencies, presumably as a consequence of reduced SERCA activity. In line with this, the hearts of adult *plna* R14del zebrafish displayed clear alternations in cardiac output. In elderly *plna* R14del mutant fish we observed cardiac remodeling, which was characterized by the accumulation of immune cells, fibroblast and fat, features reminiscent of those seen in PLN R14del patients. Importantly, our results demonstrate that drug treatment with istaroxime enhances cardiac Ca^2+^ dynamics, improves cardiac relaxation and reverses APD alternans in *pln*a R14del cardiomyocytes.

We observed a heterogeneous effect of *plna* R14del in 2-year-old zebrafish, as some fish appeared completely healthy while others displayed end-stage heart disease characterized by the accumulation of fibroblasts, immune cells and fat in the sub-epicardial region. A similar age-related heterogenous effect of the PLN R14del mutation is seen in affected patients, with fibrofatty infiltrates that are usually located in the posterolateral region of the left ventricular myocardium, right underneath the epicardium. Cardiac fibrofatty replacement in PLN R14del patients has a likely (sub)epicardial origin, which implicates that the underlying mechanisms of disease might be comparable between zebrafish and man^48^. The epicardium is a mesothelial cell layer acts as an important source of trophic signals to maintain continued growth and differentiation of the developing heart. The importance of the epicardium as a driver of morphologic changes in the heart is increasingly recognized^49^. In the adult zebrafish the epicardium responds to injury by inducing embryonic epicardial gene expression (e.g. *tbx18*) and proliferation^38,50^. After cardiac injury, the epicardial cells undergo a process called epithelial-to-mesechymal transformation (EMT). They migrate into the cardiac tissue where they can give rise to cardiac smooth muscle cells and fibroblasts^51^. Epicardial cells can also give rise to fat cells via mesenchymal transformation and peroxisome proliferator-activated receptor γ (PPARγ) activation^52^. As the remodelled tissue within *plna* R14del hearts showed strong expression of *tbx18*, it suggests that it may have an epicardial origin. This should be studied further by lineage tracing experiments.

Interestingly, *plna* R14del zebrafish presented with a remarkable higher variation in ventricular outflow peak velocity, even in absence of structural heart disease. Pulsus alternans results primarily from an alternating contractile state of the ventricle^53^. In accordance with our echocardiography measurements, we noticed a clear alternans between consecutive APs in *plna* R14del cardiomyocytes during patch clamp experiments at higher pacing frequencies. A plausible mechanism underlying these alternans is a functional change in SR Ca^2+^ pumps, resulting in alternating strong and weak beats^54,55^. Also, concomitant slow transportation of Ca^2+^ from the uptake compartment to the release compartment in the SR has been suggested as a cause of alternans^55^. Corroborating such a model, we were able to rescue the Ca^2+^ amplitudes (discussed below) and electrical alternans with istaroxime. Istaroxime is a drug which acts by releasing PLN from SERCA and thereby restores SR Ca^2+^ loading and release^43^.

In line with our presumptions, the *in vivo* Ca^2+^ transient amplitude clearly decreased in embryonic *plna*-R14del mutant fish, potentially caused by a hampered SR Ca^2+^ sequestration^56^. In contrast to earlier described observations^18^, it appears that diastolic Ca^2+^ concentrations were slightly lower in *pln*a R14del mutants. Overexpression of human PLN R14del in *Pln* null mice revealed that PLN R14del can translocate to the plasma membrane and subsequently activate NKA^11^. Increased NKA activity will lower cytosolic [Na^+^] and increase NCX mediated Ca^2+^ extrusion, thereby depleting cytosolic Ca^2+^ levels. It is tempting to speculate that this mechanism is also involved in *plna* R14del mutant fish. In line with literature, we demonstrated that istaroxime improves intracellular Ca^2+^ transient amplitude and has a positive inotropic effect on zebrafish heart function. More efficient reuptake of Ca^2+^ into the SR, by mitigated PLN-SERCA inhibition, will improve SR Ca^2+^ load, Ca^2+^ release and as a consequence elevates the Ca^2+^ transient amplitude. The enhanced Ca^2+^ transient amplitudes did not directly translate into improved contractility of the zebrafish cardiomyocyte in our study. This may be explained by high myofilament affinity for Ca^2+^, which may lead to complete myofilament saturation in both absence and presence of istaroxime. Zebrafish have the ability to adapt their myofilament Ca^2+^ sensitivity extensively in order to maintain cardiac function under a large range of temperatures (6-38°C), demanding a high Ca^2+^ buffering capacity in the cell^57^. In this study, we show that istaroxime predominantly improves the relaxation, relaxation time and filling of the zebrafish heart, rather than affecting the contractile state of the heart. The positive effect of istaroxime on cycle length relaxation time was primarily present in *plna* R14del mutants, confirming that istaroxime can improve SR Ca^2+^ storage capacity when SR Ca^2+^ sequestration is hampered^43^. Ouabain is a widely used inotropic agent in heart failure therapy. Unlike istaroxime, ouabain did not effect the Ca^2+^ transient amplitudes. This suggest a predominant lusitropic effect of istaroxime via enhanced SR Ca^2+^ sequestration, presenting istaroxime as potential effective pharmacologic agent for PLN R14del disease.

This study has also demonstrated that istaroxime shortens AP repolarization in the zebrafish heart and isolated zebrafish *pln* R14del cardiomyocytes, a phenomenon previously reported *in vitro* and *in vivo* in the chronic atrioventricular block dog model^58^. Under physiological conditions, the removal of Ca^2+^ from the cytosol is achieved via SR Ca^2+^ reuptake and via the forward mode of the NCX. Upon istaroxime exposure, the enhanced SR sequestration of Ca^2+^ through SERCA2A during diastole likely reduces the amount of Ca^2+^ extruded by the NCX and reduces its depolarizing pump current. A reduced amplitude of NCX current, in combination with enhanced Ca^2+^-induced I_Ca,L_ inactivation, therefore results in shortening of repolarization. Impaired NKA Na^+^ extrusion by istaroxime elevates intracellular Na^+^ levels and favors the reverse mode of NCX, further shortening AP repolarization. More important, in this study we show that APD alternans caused by the *plna* R14del mutation can be reversed. This highlights Ca^2+^ handling defects as underlying cause for the APD alternans in isolated ventricular *plna* R14del cardiomyocytes. APD alternans are likely not related to a prolonged repolarization, as APD_90_ did not differ between wild-type and *pln* R14del cardiomyocytes. We appreciate that the Ca^2+^ handling defects in *plna* R14del embryonic hearts and adult cardiomyocytes cannot yet be linked directly towards the structural remodeling at later stages. In PLN R14del patients the disease symptoms predominantly manifest at later ages (50-60 years of age), implying that APD alternans and impaired Ca^2+^ transient dynamics are an early phenotypical characteristic of the disease, which over time predisposes to a pathogenic Ca^2+^ handling related cardiomyopathy.

In conclusion, by introducing the R14del mutation in the endogenous zebrafish *plna* gene we generated a novel relevant zebrafish cardiomyopathy model which shows *in vivo* cardiac Ca^2+^ dysregulation and morphological features like observed in cardiac specimen from patients. Our data present possible early disease mechanisms in PLN R14del cardiomyopathy and provide usefulness of this model to explore patient specific drug treatment. Naturally, extrapolation of our findings to the human situation should be done with caution.

## Acknowledgements

We would like to thank the Hubrecht Institute animal care takers for fish care, Anko de Graaff at the imaging facility, Jeroen Korving and Harry Begthel at the histology department, and Berend de Jonge of Medical Biology (Amsterdam) for their expertise. We acknowledge the support from The Netherlands Cardio Vascular Research Initiative (CVON): the Dutch Heart Foundation, Dutch Federation of University Medical Centres, the Netherlands Organization for Health Research and Development and the Royal Netherlands Academy of Sciences (CVON-PREDICT 2012-10) and the PLN Foundation. This work was further supported by the ZonMW grant 40-00812-98-12086, the ERA-NET Cofund action N° 643578 under the European Union’s Horizon 2020 research and innovation programme and national funding organisations Canadian Institutes for Health Research (CIHR), the Netherlands Organization for Health Research and Development (ZonMw), Belgium (Flanders) Research Foundation Flanders (FWO), French National Research Agency (ANR) and E-Rare-CoHeart project.

## Author contributions

S.M.K: Study design, experiments in adult animals, data analysis and interpretation, manuscript preparation, expansion and care of the animal colony. C.J.M.vO: Study design, experiments in embryonic and adult animals, data analysis and interpretation, manuscript preparation, care of the animal colony. C.D.K: Generation of the animal model, study design, experiments in embryonic and adult animals, data analysis and interpretation, manuscript preparation, reviewing and editing, expansion and care of the animal colony. A.V: Patch clamp experiments, data interpretation and analysis, manuscript reviewing and editing. S.C.: Generation and expansion of the animal colony. Y.L.O: Cryo-sectioning and care of the animal colony. C.P. *In vivo* calcium imaging in embryonic animals. M.A.V: Funding, manuscript reviewing and editing. T.P.dB: Conceptual contribution and study design, data analysis and interpretation, manuscript reviewing and editing. T.A.B.vV: Study conceptualization, main funding and design, data analysis and interpretation, study coordination, manuscript reviewing and editing. J.B.: Study conceptualization, generation of the animal model, main funding and design, data analysis and interpretation, study coordination, manuscript reviewing and editing.

## Supplementary figures

**Figure S1.**
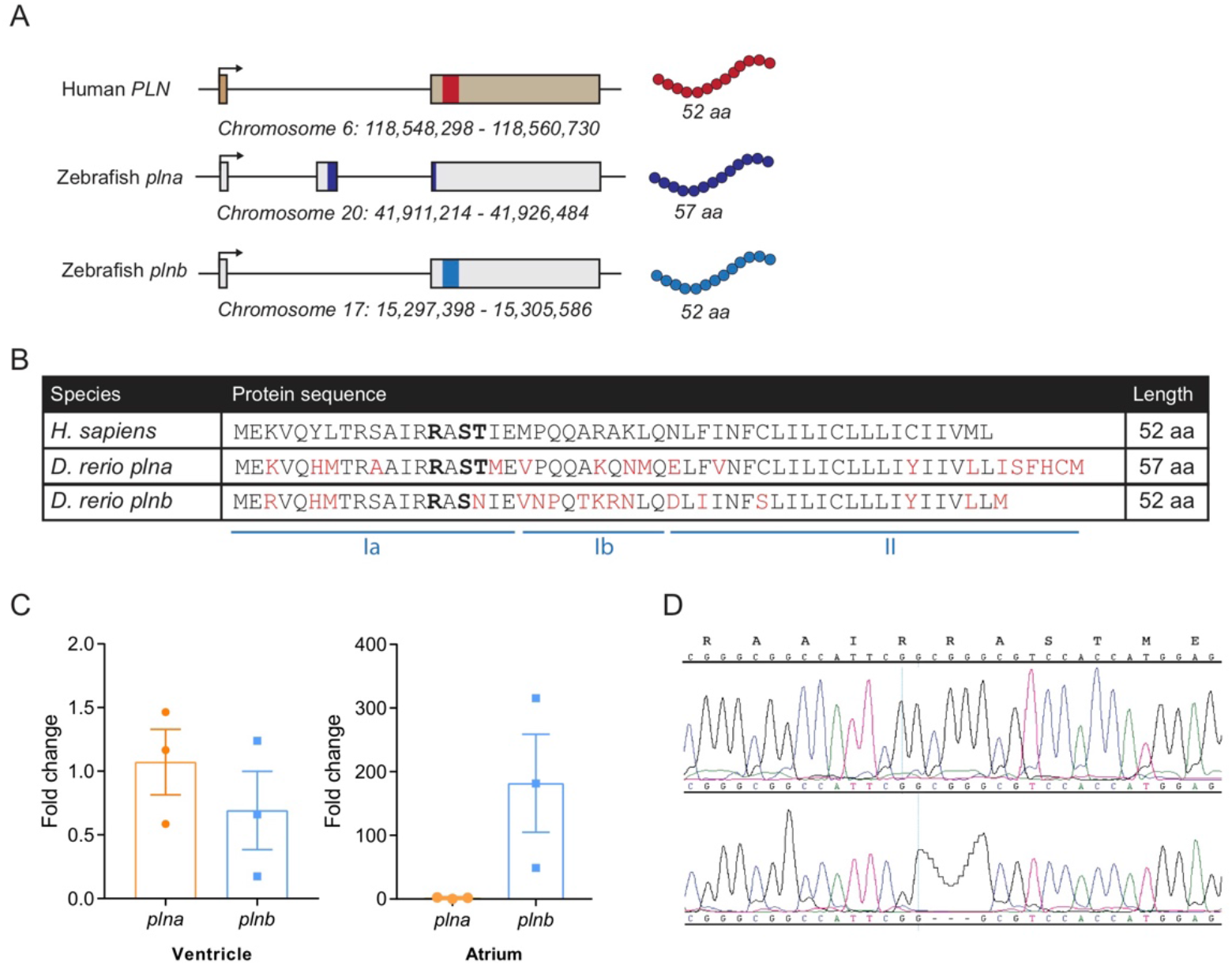
Description of zebrafish *plna* and *plnb*. **A)** In contrast to humans, zebrafish carry two *pln* genes. One gene is located on chromosome 20 (*plna*) and one on chromosome 17 (*plnb*). The R14del mutation was introduced to *plna*. **B)** Alignment of the human (*H. sapiens*) and zebrafish (*D. rerio*) protein sequences. Differences in amino acid sequence are indicated in red. The R14 and phosphorylation sites are indicated in bold. The different protein domains are indicated by la (cytosolic), lb (linker), and ll (transmembrane). **C)** Quantitative PCR plot shows the expression levels of *plna* and *plnb* in ventricle (n=3) and atrium (n=3) of wild-type fish. **D)** Sequencing peaks of *plna* showing the R14del mutation from a wild-type and a *plna* R14del mutant fish. In the R14del fish, the Arginine on amino acid location 14 has evidently been deleted. aa: amino acids, Neg.: negative.

**Figure S2.**
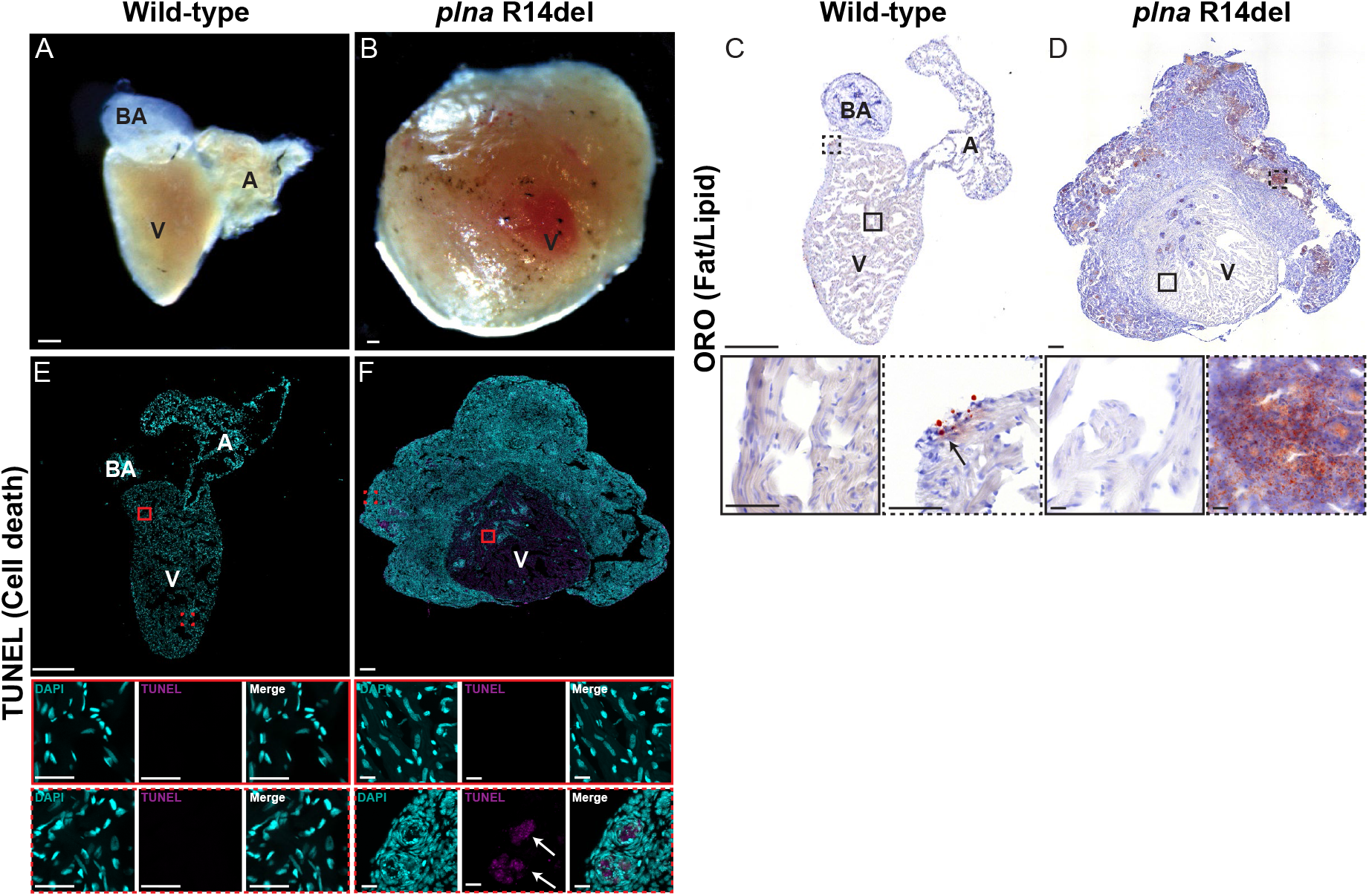
Fibrofatty replacement and cell death in adult *plna* R14del hearts. **A-B)** Bright field images of isolated wild-type and *plna* R14del mutant zebrafish hearts (10 months). **C-D)** Oil Red O staining on the hearts of the two genotypes is shown, with 40x zoom-in on the myocardium and ORO stained region. **E-F)** TUNEL fluorescent staining to indicate apoptosis, where DAPI+ nuclei are shown in cyan and TUNEL+ nuclei in magenta. Images were taken at a zoom of 20x for whole heart and at a zoom of 63x for TUNEL staining. Scale bars are 200μm for whole heart tile scans and 20μm or 50 μm for zoom-in regions at 40x and 63x, respectively. A: atrium, V: ventricle, BA: bulbus arteriosus.

**Figure S3.**
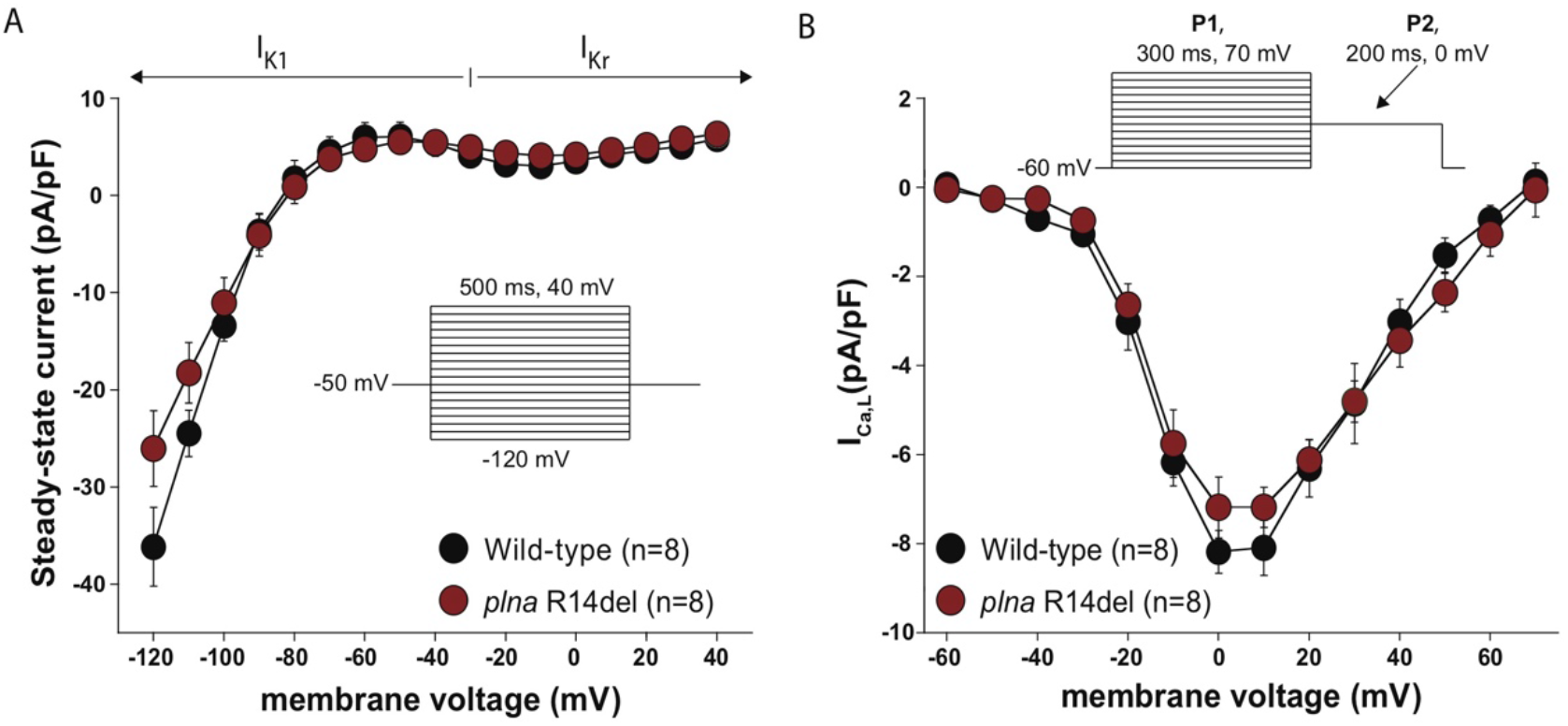
Effect of the *plna* R14del mutation on membrane currents in isolated cardiomyocytes. **A)** Current-voltage (I-V) relationships of I_K1_ and I_Kr_ in cardiomyocytes from wild-type (WT) and *plna* R14del mutant zebrafish (mean ± SEM, unpaired Students t-test). Inset, voltage clamp protocol used. **B)** I-V relationships of I_Ca,L_ in WT and *plna* R14del zebrafish cardiomyocytes (mean ± SEM, unpaired Students t-test). Inset, voltage clamp protocol used. WT: wild type, ms: milliseconds, mV: millivolt, I_K1_: inward rectifier potassium current, I_Kr_: rapidly activated delayed rectifier potassium current, I_Ca,L_: L-type calcium current.

**Figure S4.**
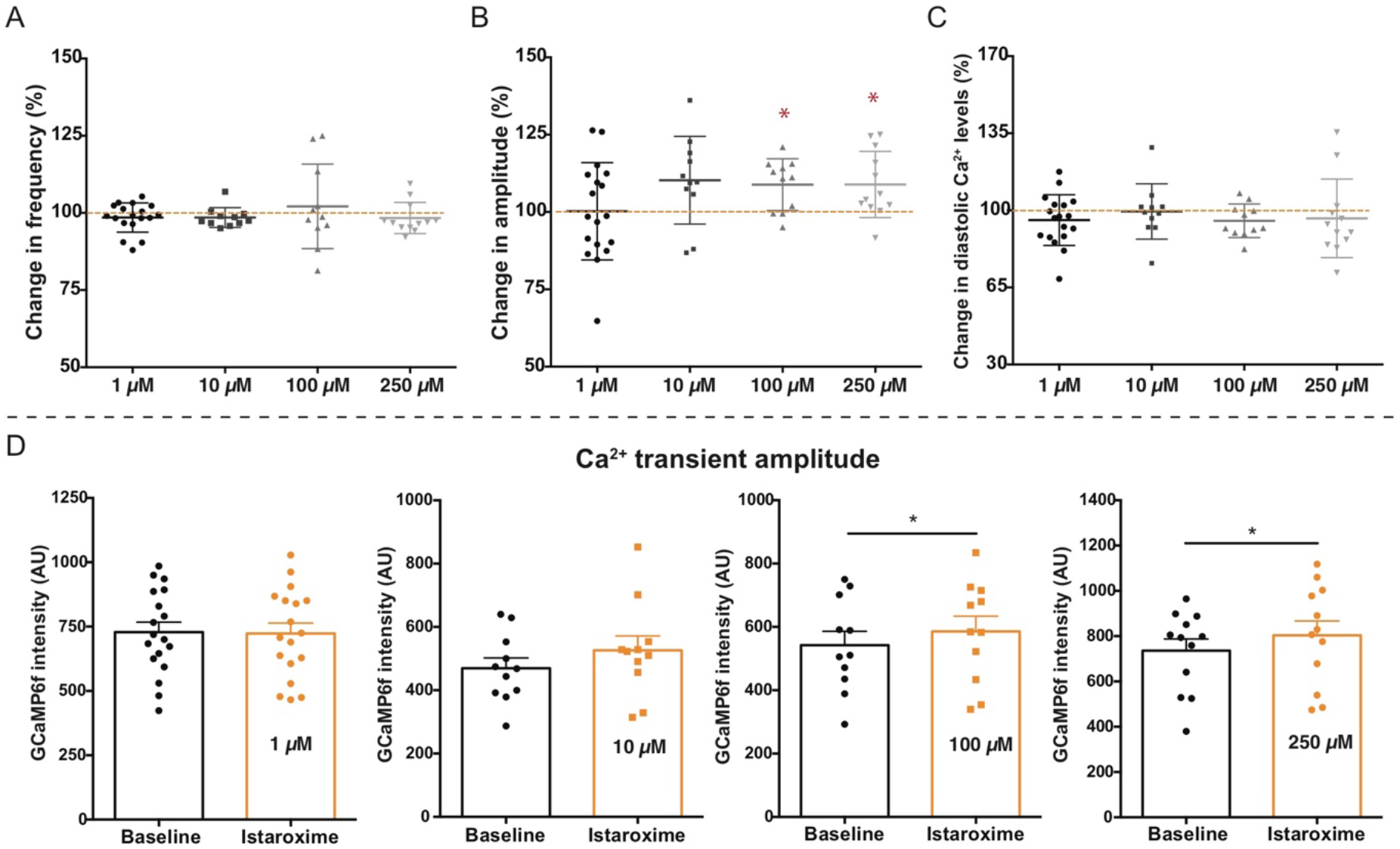
Dose response effect of istaroxime in GCaMP6f embryos. **A-C)** Effect of increasing concentrations of istaroxime (1-250 μM) on calcium (Ca^2+^) transient parameters in GCaMP6f zebrafish. Values are presented as relative change (in %) from grouped baseline values of all concentrations. Parameters include **A)** the frequency of Ca^2+^ transients, **B)** the Ca^2+^ transient amplitude and **C**) diastolic Ca^2+^ levels (mean ± STDEV, *p≤0.05, baseline n=51, 1 μM n= 17, 10 μM n= 11, 100 μM n= 11, 250 μM n= 12, unpaired Students t-test). **D**) The effect of different istaroxime concentrations (1-250 μM) on the Ca^2+^ transient amplitude in GCaMP6f zebrafish. Bar graphs represent baseline values and values after incubation with istaroxime (mean ± SEM, *p≤0.05, 1 μM n= 18, 10 μM n= 11, 100 μM n= 11, 250 μM n= 12, paired Students t-test). AU: arbitrary units.

**Figure S5.**
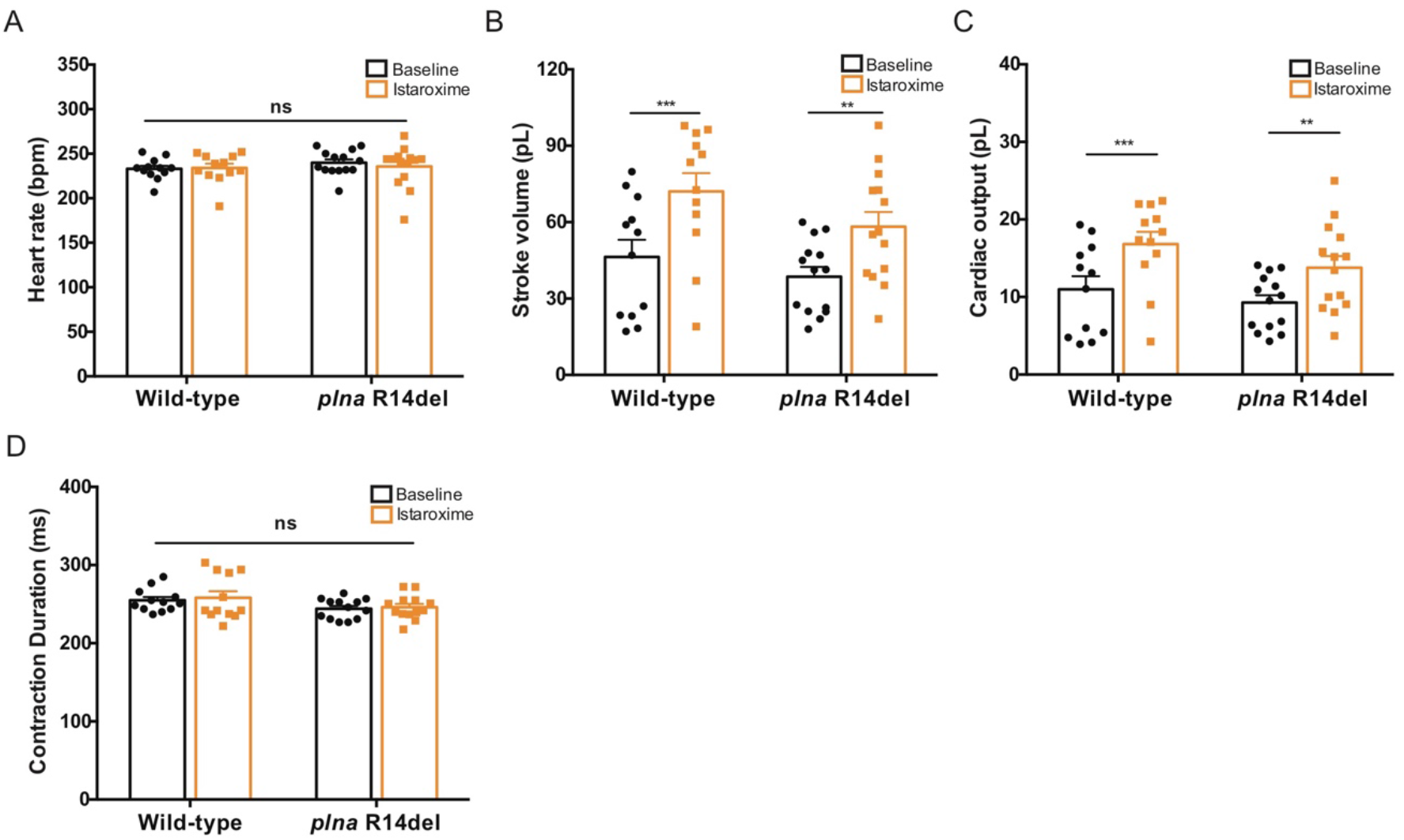
Effect of istaroxime on cardiac contractility parameters in wild-type and *plna* R14del mutants. **A-D)** Cardiac contractility parameters examined in wild-type (WT) and *plna* R14del mutant embryos at baseline and after incubation with 100 μM istaroxime (mean ± SEM, ** p≤0.01, *** p≤0.001, WT baseline n=12, WT istaroxime n=12, *plna* baseline n=14, *plna* istaroxime n=14, paired Students t-test), including heart rate (**A**), stroke volume (**B**), cardiac output (**C**) and contractile cycle length contraction duration (**D**). bpm: beats per minute, pL: Pico litre, ms: milliseconds.

**Figure S6.**
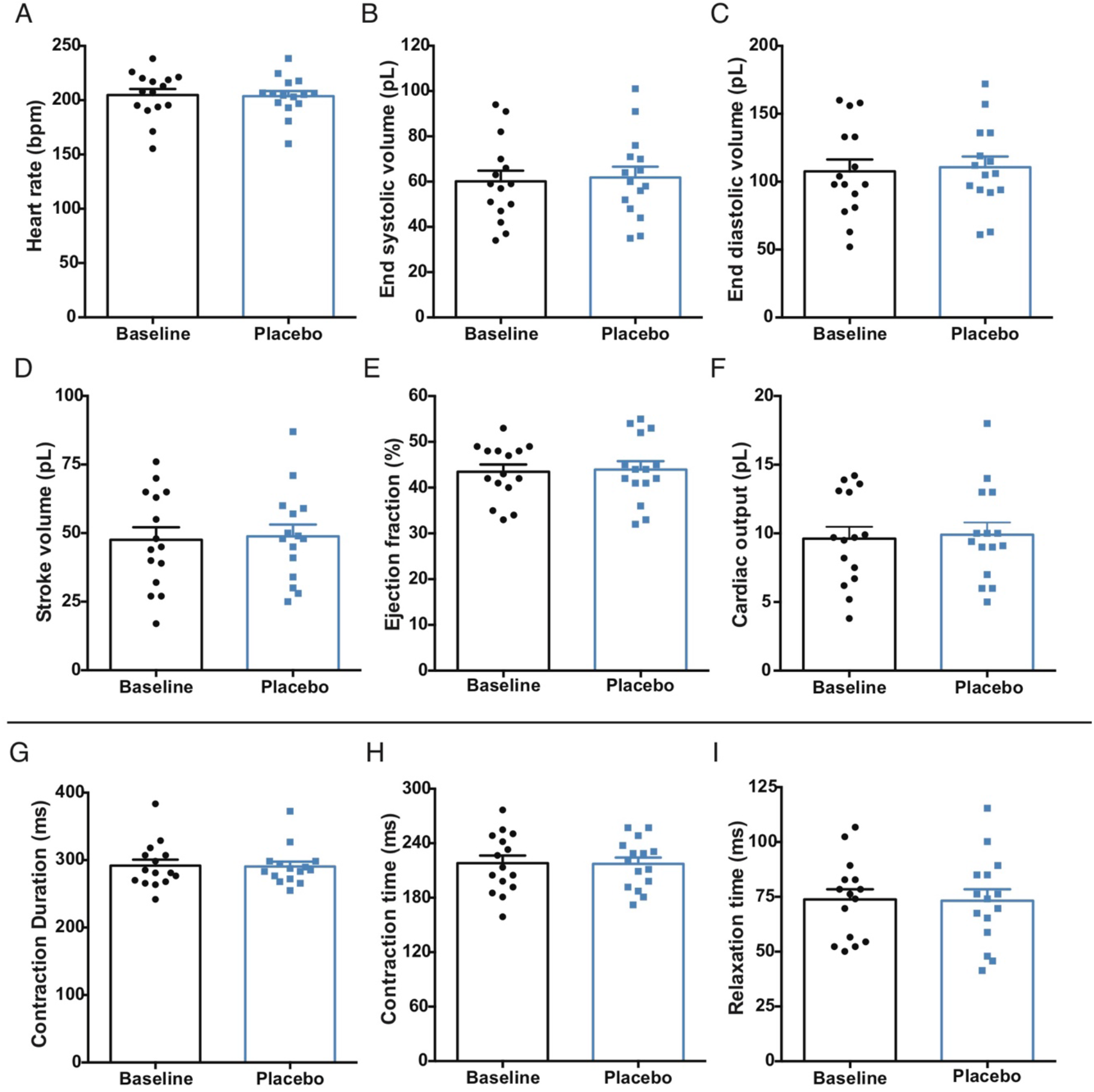
Effect of placebo treatment on cardiac contractility parameters in wild-type embryos. **A-I)** Cardiac contractility parameters examined in wild-type (WT) embryos at baseline, and 30 minutes after replacing the incubation medium (E3 water + MS222) with the exact similar medium (mean ± SEM, baseline n=15, placebo n=15, paired Students t-test), including heart rate (**A**), end systolic volume (**B**), end diastolic volume (**C**), stroke volume (**D**), ejection fraction (**E)**, cardiac output (**F**), contractile cycle length contraction duration (**G**), contractile cycle length contraction time (**H**) and contractile cycle length relaxation time (**I**). bpm: beats per minute, pL: Pico litre, ms: milliseconds.

**Figure S7.**
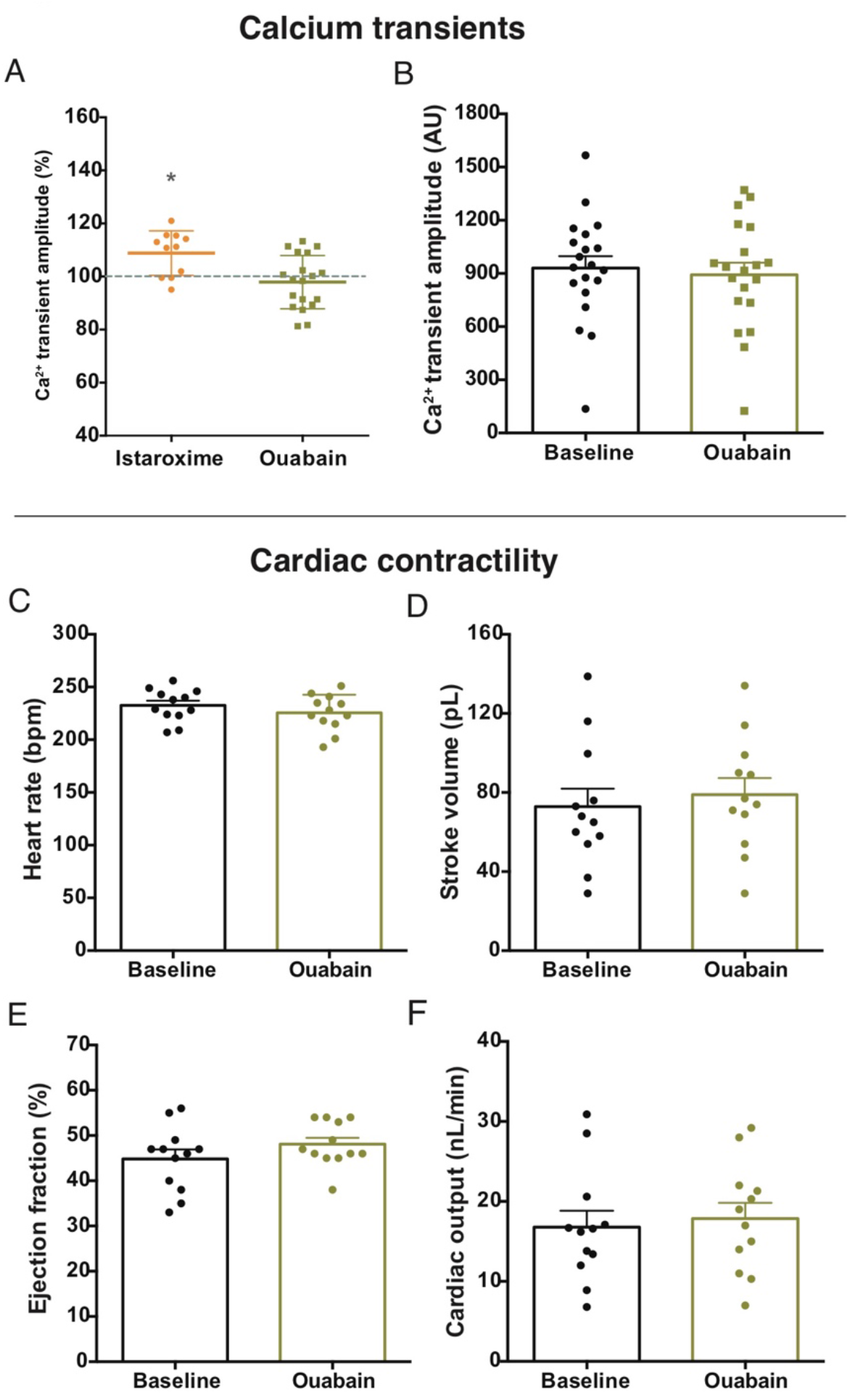
In vivo effect of ouabain on calcium transient amplitude and cardiac contractility in GCaMP6f embryos. **A)** Effect of 100 μM istaroxime and 100 μM ouabain on the calcium (Ca^2+^) transient amplitude in GCaMP6f zebrafish. Values are presented as relative change (in %) from baseline values (mean ± SEM, * p≤0.05, baseline n=35, istaroxime n=15, ouabain n=20, paired Students t-test). **B)** The Ca^2+^ transient amplitude at baseline and after incubation with 100 μM ouabain (mean ± SEM, n=20, paired Students t-test)**. C-F)** Cardiac contractility parameters examined in GCaMP6f embryonic zebrafish at baseline and after incubation with 100 μM ouabain (mean ± SEM, baseline n=12, ouabain n=12, paired Students t-test), including heart rate (**C**), stroke volume (**D**), ejection fraction (**E**) and cardiac output (**F**). bpm: beats per minute, pL: Pico litre, nL/min: nanolitre per minute

**Figure S8.**
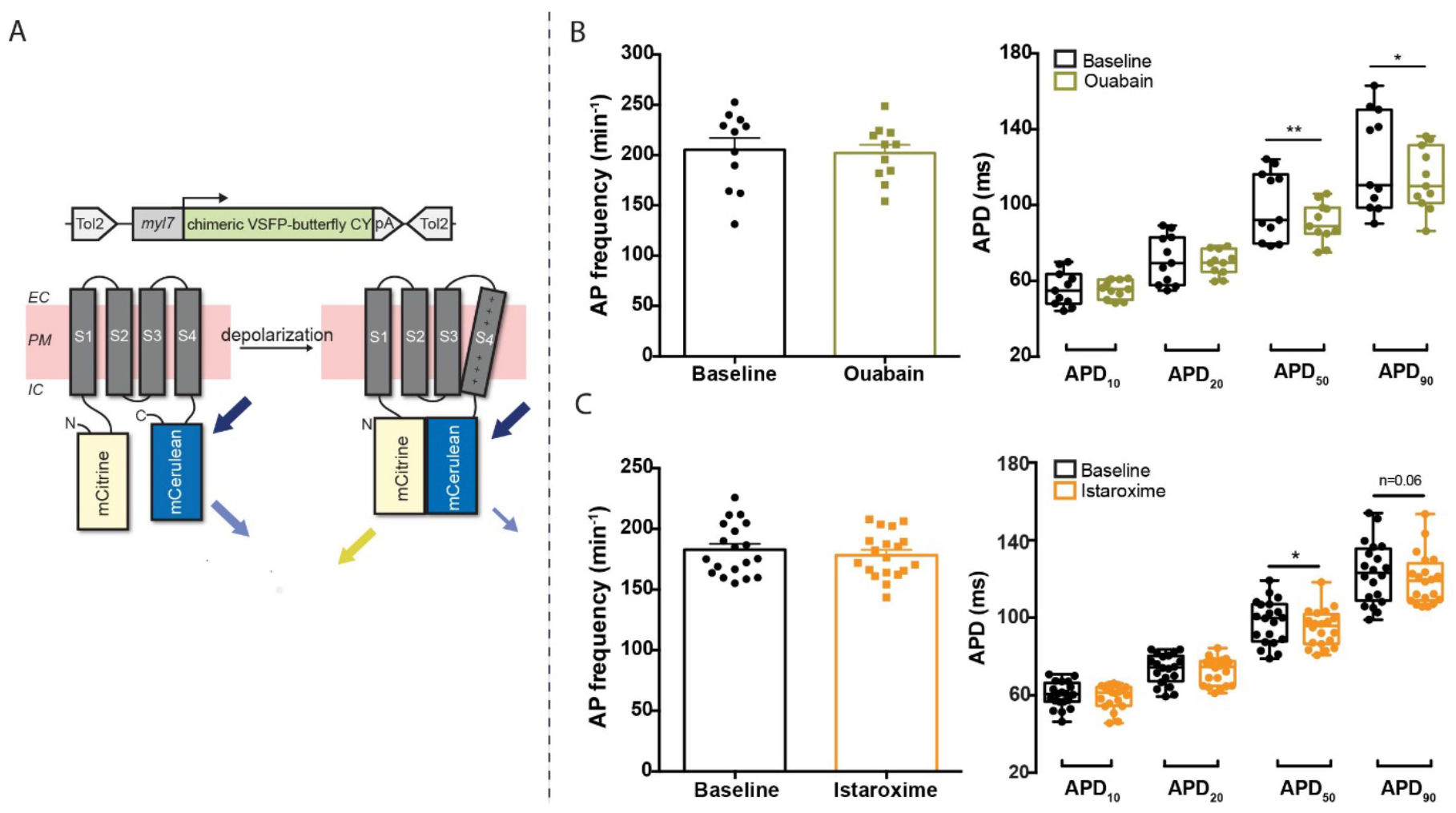
In vivo effect of istaroxime and ouabain on action potential parameters in VSFP-butterfly CY embryos. **A)** DNA construct and concept of the sensing mechanism of chimeric voltage sensitive fluorescent protein (VSFP)-butterfly CY. Chimeric VSFP-butterfly CY was placed under control of the *myl7* promoter to restrict its expression to the heart. The senor consists of a voltage sensitive domain with transmembrane segments S1-S4, sandwiched between a fluorescence resonance energy transfer (FRET) pair of the fluorescent proteins mCitrine and mCerulean. Movement of S4 upon membrane depolarization translates into a change of FRET efficiency. **B)** Frequency of action potentials and APD parameters at baseline and after incubation with 100 μM ouabain. (mean ± SEM, * p≤0.05, ** p≤0.01, n=11, paired Students t-test). **C)** Frequency of action potentials and APD parameters at baseline and after incubation with 100 μM istaroxime (mean ± SEM, * p≤0.05, n=19, paired Students t-test). min^−1^: per minute, ms: milliseconds, APD_10_: AP duration at 10% of repolarization, APD_20_: AP duration at 20% of repolarization, APD_50_: AP duration at 50% of repolarization, APD_90_: AP duration at 90% of repolarization.

**Table S1.** Echocardiographic measurements of 6 and 10 months old wild-type (WT) and *plna* R14del zebrafish. Values are mean ± SEM. VOT; ventricular outflow tract, PV; peak velocity, bpm; beats per minute, mm; millimeters, s; second, pL; Pico litre, mL/min, milliliter per minute. * p≤0.05, ** p≤0.01, *** p≤0.001, **** p≤0.0001. * 6 months old fish versus 10 months old fish, * WT versus plna R14del. Two-way ANOVA

| Echocardiographic parameters |  |  |  |  |  |  |  |  |
| --- | --- | --- | --- | --- | --- | --- | --- | --- |
|  | n | Heart rate (bpm) | VOT diameter (mm) | VOT surface (mm<sup>2</sup>) | VOT PV (mm/s) | Stroke volume (pL) | Cardiac output (mL/min) | VOT PV variation (mm/s) |
| Wildtype 6 Months | 10 | 92.39 ± 4.08 | 0.379 ± 0.014 | 0.203 ± 0.028 | 100.79 ± 11.50 | 2.88 ± 0.476 | 0.236 ± 0.049 | 4.43 ± 0.46 |
| *plna* R14del 6 Months | 10 | 99.75 ± 5.29 | 0.368 ± 0.021 | 0.219 ± 0.025 | 101.64 ± 10.33 | 2.65 ± 0.53 | 0.253 ± 0.051 | 5.78 ± 0.84 |
| Wildtype 10 Months | 21 | 98.97 ± 4.93 | 0.440 ± 0.009 ** | 0.306 ± 0.011 ** | 146.31 ± 6.57* | 5.74 ± 0.41** | 0.554 ± 0.042** | 2.97 ± 0.332 |
| *plna* R14del 10 Months | 21 | 93.93 ± 3.92 | 0.446 ± 0.008 *** | 0.314 ± 0.010 *** | 159.07 ± 10.20** | 5.99 ± 0.40**** | 0.552 ± 0.038*** | 6.30 ± 0.85* |

\*p 6 months vs. 10 Months
\*p WT vs. *plna* R14del 10 Months

## References

1. Corrado, D., Basso, C. & Judge, D. P. Arrhythmogenic Cardiomyopathy. Circ. Res. 121, 784–802 (2017).

2. Podgoršek, B., Poglajen, G., Cerar, A., Šinkovec, M. & Vrtovec, B. Arrhythmogenic cardiomyopathy. Zdr. Vestn. 87, 599–617 (2019).

3. Sommariva, E., Stadiotti, I., Perrucci, G. L., Tondo, C. & Pompilio, G. Cell models of arrhythmogenic cardiomyopathy: Advances and opportunities. DMM Disease Models and Mechanisms 10, 823–835 (2017).

4. Hof, I. E. et al. Prevalence and cardiac phenotype of patients with a phospholamban mutation. Netherlands Hear. J. 27, 64–69 (2019).

5. DeWitt, M. M., MacLeod, H. M., Soliven, B. & McNally, E. M. Phospholamban R14 Deletion Results in Late-Onset, Mild, Hereditary Dilated Cardiomyopathy. J. Am. Coll. Cardiol. 48, 1396–1398 (2006).

6. Haghighi, K. et al. A mutation in the human phospholamban gene, deleting arginine 14, results in lethal, hereditary cardiomyopathy. Proc. Natl. Acad. Sci. 103, 1388–1393 (2006).

7. López-Ayala, J. M. et al. Phospholamban p.arg14del Mutation in a Spanish Family With Arrhythmogenic Cardiomyopathy: Evidence for a European Founder Mutation. Rev. Española Cardiol. (English Ed. 68, 346–349 (2015).

8. Posch, M. G. et al. Genetic deletion of arginine 14 in phospholamban causes dilated cardiomyopathy with attenuated electrocardiographic R amplitudes. Hear. Rhythm 6, 480–486 (2009).

9. te Rijdt, W. P. et al. Phospholamban immunostaining is a highly sensitive and specific method for diagnosing phospholamban p.Arg14del cardiomyopathy. Cardiovasc. Pathol. 30, 23–26 (2017).

10. Van Der Zwaag, P. A. et al. Recurrent and founder mutations in the Netherlands - Phospholamban p.Arg14del mutation causes arrhythmogenic cardiomyopathy. Found. Mutat. Inherit. Card. Dis. Netherlands 81–87 (2014). doi:10.1007/978-90-368-0705-0_11

11. Haghighi, K. et al. The human phospholamban Arg14-deletion mutant localizes to plasma membrane and interacts with the Na/K-ATPase. J. Mol. Cell. Cardiol. 52, 773–82 (2012).

12. Van Rijsingen, I. A. W. et al. Outcome in phospholamban R14del carriers results of a large multicentre cohort study. Circ. Cardiovasc. Genet. 7, 455–465 (2014).

13. MacLennan, D. H. & Kranias, E. G. Phospholamban: A crucial regulator of cardiac contractility. Nat. Rev. Mol. Cell Biol. 4, 566–577 (2003).

14. Li, L. I., Chu, G., Kranias, E. G., Bers, D. M. & Li, L. Downloaded from physiology.org/journal/ajpheart at Utrecht Univ Lib (143.121.196.015) on. (2019).

15. Mazzocchi, G. et al. Phospholamban ablation rescues the enhanced propensity to arrhythmias of mice with CaMKII-constitutive phosphorylation of RyR2 at site S2814. J. Physiol. 594, 3005–3030 (2016).

16. Del Monte, F., Harding, S. E., Dec, G. W., Gwathmey, J. K. & Hajjar, R. J. Targeting Phospholamban by Gene Transfer in Human Heart Failure.

17. Vostrikov, V. V., Soller, K. J., Ha, K. N., Gopinath, T. & Veglia, G. Effects of naturally occurring arginine 14 deletion on phospholamban conformational dynamics and membrane interactions. Biochim. Biophys. Acta - Biomembr. 1848, 315–322 (2015).

18. Karakikes, I. et al. Correction of human phospholamban R14del mutation associated with cardiomyopathy using targeted nucleases and combination therapy. Nat. Commun. 6, 6955 (2015).

19. Lieschke, G. J. & Currie, P. D. Animal models of human disease: Zebrafish swim into view. Nature Reviews Genetics 8, 353–367 (2007).

20. Poon, K. L. & Brand, T. The zebrafish model system in cardiovascular research: A tiny fish with mighty prospects. Glob. Cardiol. Sci. Pract. 2013, 4 (2013).

21. Howe, K. et al. The zebrafish reference genome sequence and its relationship to the human genome. Nature 496, 498–503 (2013).

22. MacRae, C. A. & Peterson, R. T. Zebrafish as tools for drug discovery. Nature Reviews Drug Discovery 14, 721–731 (2015).

23. Prykhozhij, S. V. & Berman, J. N. Zebrafish knock-ins swim into the mainstream. DMM Disease Models and Mechanisms 11, (2018).

24. Tessadori, F. et al. Effective CRISPR/Cas9-based nucleotide editing in zebrafish to model human genetic cardiovascular disorders. Dis. Model. Mech. 11, dmm035469 (2018).

25. Farr, G. H., Imani, K., Pouv, D. & Maves, L. Functional testing of a human PBX3 variant in zebrafish reveals a potential modifier role in congenital heart defects. DMM Dis. Model. Mech. 11, (2018).

26. Westerfield, M. The Zebrafish Book: A Guide for the Laboratory Use of Zebrafish (Danio Rerio). (University of Oregon Press, 2000).

27. Moorman, A. F. M., Houweling, A. C., De Boer, P. A. J. & Christoffels, V. M. Sensitive Nonradioactive Detection of mRNA in Tissue Sections: Novel Application of the Whole-mount In Situ Hybridization Protocol. The Journal of Histochemistry & Cytochemistry 49, (2001).

28. Yelon, D., Horne, S. A. & Stainier, D. Y. R. Restricted expression of cardiac myosin genes reveals regulated aspects of heart tube assembly in zebrafish. Dev. Biol. 214, 23–37 (1999).

29. Begemann, G., Gibert, Y., Meyer, A. & Ingham, P. W. Cloning of zebrafish T-box genes tbx15 and tbx18 and their expression during embryonic development. Mech. Dev. 114, 137–141 (2002).

30. Poss, K. D., Wilson, L. G. & Keating, M. T. Heart Regeneration in Zebrafish. Science 298, (2002).

31. Bancroft, J. D. & Stevens, A. Theory and Practice of Histological Techniques. (World Publishing Corporation, 1991).

32. Wang, L. W. et al. Standardized echocardiographic assessment of cardiac function in normal adult zebrafish and heart disease models. Dis. Model. Mech. 10, 63–76 (2017).

33. Tessadori, F. et al. Identification and Functional Characterization of Cardiac Pacemaker Cells in Zebrafish. PLoS One 7, (2012).

34. Barry, P. H. & Lynch, J. W. Liquid junction potentials and small cell effects in patch-clamp analysis. J. Membr. Biol. 121, 101–117 (1991).

35. van Opbergen, C. J. M. et al. Optogenetic sensors in the zebrafish heart: a novel in vivo electrophysiological tool to study cardiac arrhythmogenesis. Theranostics 8, 4750–4764 (2018).

36. White, R. M. et al. Transparent Adult Zebrafish as a Tool for In Vivo Transplantation Analysis. Cell Stem Cell 2, 183–189 (2008).

37. Livak, K. J. & Schmittgen, T. D. Analysis of relative gene expression data using real-time quantitative PCR and the 2-ΔΔCT method. Methods 25, 402–408 (2001).

38. Lepilina, A. et al. A Dynamic Epicardial Injury Response Supports Progenitor Cell Activity during Zebrafish Heart Regeneration. Cell 127, 607–619 (2006).

39. Talman, V. & Ruskoaho, H. Cardiac fibrosis in myocardial infarction-from repair and remodeling to regeneration. Cell Tissue Res. 365, 563–81 (2016).

40. Edwards, J. N. & Blatter, L. A. Cardiac alternans and intracellular calcium cycling. Clin. Exp. Pharmacol. Physiol. 41, 524–32 (2014).

41. Verkerk, A. O. & Remme, C. A. Zebrafish: a novel research tool for cardiac (patho)electrophysiology and ion channel disorders. Front. Physiol. 3, 255 (2012).

42. Kanaporis, G. & Blatter, L. A. The Mechanisms of Calcium Cycling and Action Potential Dynamics in Cardiac Alternans. Circ. Res. 116, 846–856 (2015).

43. Ferrandi, M. et al. Istaroxime stimulates SERCA2a and accelerates calcium cycling in heart failure by relieving phospholamban inhibition. Br. J. Pharmacol. 169, 1849–1861 (2013).

44. Gobbini, M. et al. Novel analogues of Istaroxime, a potent inhibitor of Na+,K+-ATPase: Synthesis, structure-activity relationship and 3D-quantitative structure-activity relationship of derivatives at position 6 on the androstane scaffold. Bioorganic Med. Chem. 18, 4275–4299 (2010).

45. Alevizopoulos, K. et al. Functional characterization and anti-cancer action of the clinical phase II cardiac Na+/K+ ATPase inhibitor istaroxime: in vitro and in vivo properties and cross talk with the membrane androgen receptor. Oncotarget 7, 24415–28 (2016).

46. te Rijdt, W. P. et al. Myocardial fibrosis as an early feature in phospholamban p.Arg14del mutation carriers: phenotypic insights from cardiovascular magnetic resonance imaging. Eur. Hear. J. - Cardiovasc. Imaging 20, 92–100 (2019).

47. Aditya, S. & Rattan, A. Istaroxime: A rising star in acute heart failure. J. Pharmacol. Pharmacother. 3, 353 (2012).

48. Sepehrkhouy, S. et al. Distinct fibrosis pattern in desmosomal and phospholamban mutation carriers in hereditary cardiomyopathies. Hear. Rhythm 14, 1024–1032 (2017).

49. Smart, N. & Riley, P. R. The epicardium as a candidate for heart regeneration. Future Cardiol. 8, 53–69 (2012).

50. Torres, M., Mercader, N., Peralta, M., Martin, V. & Gonzalez-Rosa, J. M. Extensive scar formation and regression during heart regeneration after cryoinjury in zebrafish. Development 138, 1663–1674 (2011).

51. Sánchez-Iranzo, H. et al. Transient fibrosis resolves via fibroblast inactivation in the regenerating zebrafish heart. Proc. Natl. Acad. Sci. 115, 4188–4193 (2018).

52. Yamaguchi, Y. et al. Adipogenesis and epicardial adipose tissue: A novel fate of the epicardium induced by mesenchymal transformation and PPARγ activation. Proc. Natl. Acad. Sci. 112, 2070–2075 (2015).

53. Nguyen, T., Cao, L.-B., Tran, M. & Movahed, A. Biventricular pulsus alternans: An echocardiographic finding in patient with pulmonary embolism. World J. Clin. cases 1, 162–5 (2013).

54. Schmidt, A. G. et al. Cardiac-specific Overexpression of Calsequestrin Results in Left Ventricular Hypertrophy, Depressed Force–frequency Relation and Pulsus Alternans In Vivo. J. Mol. Cell. Cardiol. 32, 1735–1744 (2000).

55. Kihara, Y. & Morgan, J. P. Abnormal Cai2+ handling is the primary cause of mechanical alternans: study in ferret ventricular muscles. Am. J. Physiol. Circ. Physiol. 261, H1746–H1755 (1991).

56. Haghighi, K. et al. A mutation in the human phospholamban gene, deleting arginine 14, results in lethal, hereditary cardiomyopathy. Proc. Natl. Acad. Sci. U. S. A. 103, 1388–93 (2006).

57. Stevens, C. M. et al. Characterization of Zebrafish Cardiac and Slow Skeletal Troponin C Paralogs by MD Simulation and ITC. Biophys. J. 111, 38–49 (2016).

58. Bossu, A. et al. Istaroxime, a positive inotropic agent devoid of proarrhythmic properties in sensitive chronic atrioventricular block dogs. Pharmacol. Res. 133, 132–140 (2018).

